# Molecular mechanism of the pH-dependent calcium affinity in langerin

**DOI:** 10.1101/2020.03.11.986851

**Authors:** Jan-O. Joswig, Jennifer Anders, Hengxi Zhang, Christoph Rademacher, Bettina G. Keller

**Affiliations:** Department of Biology, Chemistry, and Pharmacy, Freie Universität Berlin, Arnimallee 22, 14195 Berlin, Germany; Department of Biomolecular Systems, Max Planck Institute of Colloids and Interfaces, Am Mühlenberg 1, 14424 Potsdam, Germany; Department of Pharmaceutical Chemistry, University of Vienna, Althanstraße 14, 1090 Vienna, Austria; Department of Microbiology and Immunobiology, Max F. Perutz Laboratories, University of Vienna, Campus Vienna Biocenter 5, 1030 Vienna, Austria

**Keywords:** pH regulation, allosteric regulation, conformational change, calcium-binding protein, pattern recognition receptor (PRR), molecular dynamics, computer modeling

## Abstract

The C-type lectin receptor langerin plays a vital role in the mammalian defense against invading pathogens. Its function hinges on the affinity to its co-factor Ca^2+^ which in turn is regulated by the pH. We studied the structural consequences of protonating the allosteric pH-sensor histidine H294 by molecular dynamics simulations (total simulation time: about 120 µs) and Markov models. We discovered a mechanism in which the signal that the pH has dropped is transferred to the Ca^2+^-binding site without transferring the initial proton. Instead, protonation of H294 unlocks a conformation in which a protonated lysine side-chain forms a hydrogen bond with a Ca^2+^-coordinating aspartic acid. This destabilizes Ca^2+^ in the binding pocket, which we probed by steered molecular dynamics. After Ca^2+^-release, the proton is likely transferred to the aspartic acid and stabilized by a dyad with a nearby glutamic acid, triggering a conformational transition and thus preventing Ca^2+^-rebinding.

## Introduction

When pathogens invade a mammal (or more specifically: a human), Langerhans cells capture some of the pathogens, process them, and present antigens to the adaptive immune system. The swift activation of the adaptive immune system is critical for the survival of the mammal, and langerin plays a vital role in this process. Langerin is a transmembrane carbohydrate receptor, which is expressed by Langerhans cells of mammalian skin and mucosa (1, 2). It belongs to the class of type II C-type lectin receptors (3, 4). It detects pathogens such as influenza virus (5), measles virus (6), HIV (7), fungi (8), mycobacteria (9), and bacteria (10).

Langerin recognizes these pathogens by binding to carbohydrates on the pathogen surface. Its carbohydrate binding pocket contains a Ca^2+^-cation as co-factor that is essential for carbohydrate binding, and thus for the capture of pathogens. After the initial binding event, the pathogen is captured in an endocytic vesicle, and langerin releases the pathogen into the endosome (Fig. 1a) (1, 2, 7, 11). This cargo release is triggered by a drop of pH from 7 in the extracellular medium to 5.5 to 6 in the early endosome (12) and by a substantial drop in the Ca^2+^-concentration from about 1 to 2 mM to a value in the micro molar range (13–15).

**Figure 1.**
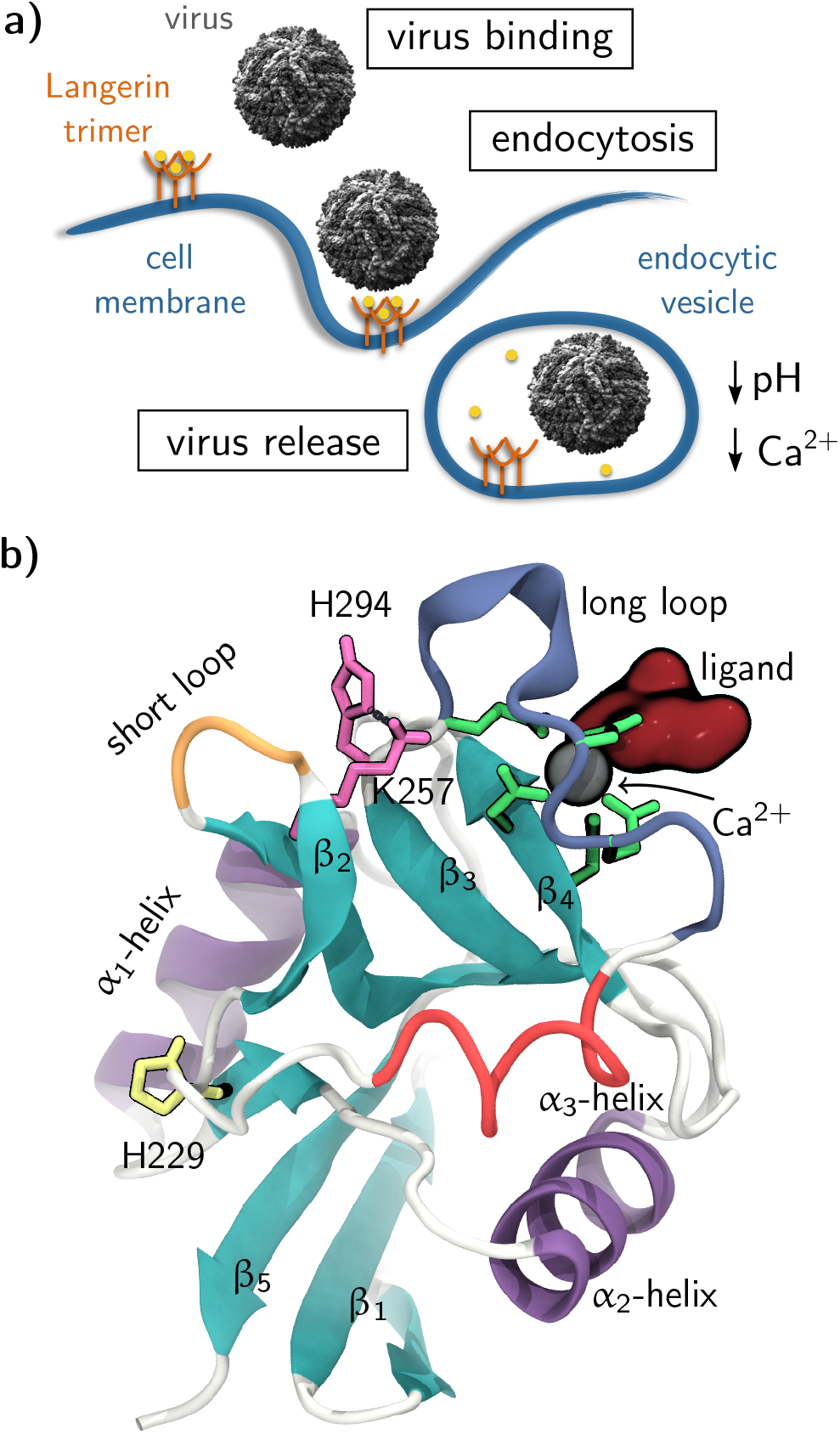
**a)** Langerin’s function as an endocytic pattern recognition receptor (16). **b)** Langerin carbohydrate recognition domain (PDB-ID 3p5g (17)).

The pH-dependent cargo-release is accomplished by a fascinating mechanism in which various chemical equilibria are carefully balanced. To be able to release the cargo into the more acidic endosome, the carbohydrate affinity of langerin needs to be pH-dependent. However, the change in pH does not affect the carbohydrate binding itself. Instead langerin depends on a Ca^2+^ co-factor for carbohydrate binding, and the observed pH-dependence of the carbohydrate affinity is caused by an underlying pH-dependence of the Ca^2+^- affinity (18). We previously showed that the Ca^2+^-affinity is lower at pH 6 than at pH 7. The pH-sensitivity, measured as the difference in the Ca^2+^-binding free-energies, is ΔΔ*G* = 5.1 kJ mol^−1^ (18). At high Ca^2+^-concentrations (10 mM) the carbohydrate affinity ceases to be pH-dependent, because the excess in Ca^2+^-ions outweighs any change in Ca^2+^-affinity due to a change in pH. However, in the endosome the Ca^2+^-concentration is low. Thus, the drop in pH from the extracellular medium to the endosome causes a decrease in Ca^2+^-affinity, and the unbinding of the Ca^2+^ co-factor leads to the dissociation of the carbohydrate ligand and to the release of the pathogen. Similar, pH-sensitivities of either ligand or Ca^2+^-affinities have been observed for several other C-type lectins (19), including ASGPR (14, 20, 21) the macrophage mannose receptor (22), DC-SIGN and DC-SIGNR (23–26), and LSECtin (27) (example structures in SI Fig. 32). In DC-SIGNR and LSECtin, which have a different biological role than langerin, a drop in pH causes an increase in ligand affinity. The mechanisms underlying the regulation by the pH in C-type lectins are highly diverse and not yet studied in detail.

The observation that the Ca^2+^-affinity in C-type lectins is pH-dependent is surprising. First, when a carbohydrate (and attached to it an entire virus) is bound to a C-type lectin, the Ca^2+^-binding site is almost certainly not solvent exposed. Second, the Ca^2+^-ion in C-type lectins is coordinated by either aspartate or glutamate side-chains, whose reference p*K*_a_-values (28) (in water at 25 °C) are 3.71 (aspartate) and 4.15 (glutamate). By themselves, these residues are not sensitive to a change in pH from 7 to 6. Pairs of acidic residues can in principle form a protonated dyad, which is the close arrangement of two residues with acidic side-chains such that protonation of their carboxyl groups is coupled. This results in an increased p*K*_a_ of the protonated residue, stabilized by the unprotonated form of the other group. Prominent examples of this effect are found in the proteins HIV-1 protease (29, 30), BACE-1 (31) BACE-2, and CatD, where it can increase the p*K*_a_ of aspartic acid from its reference value to 5.2 (32). The presence of organic ligands can increase these values further (33). However, a protonated dyad can only form if Ca^2+^ has already left the binding pocket. So the question arises: how do C-type lectins sense a change in pH, and how does this lead to the release of Ca^2+^?

For langerin we previously identified the histidine residue H294 as a partial pH sensor that regulates the Ca^2+^-affinity (18). The reference p*K*_a_ of histidine is 6.04 (in water at 25 °C) (28) which makes it sensitive to a pH change from 7 to 6. When H294 is mutated to A294, the pH-sensitivity is about 40 % smaller than in the wild-type (ΔΔ*G* = 3.1 kJ mol^−1^ upon a change in pH from 7 to 6). Because the histidine side-chain points away from the Ca^2+^- binding site, it is unlikely that the decrease in Ca^2+^-affinity is caused by electrostatic repulsion between the protonated histidine and the Ca^2+^-cation. This mechanism has been suggested for the C-type lectin ASGPR, in which however the histidine pH-sensor is located directly underneath the Ca^2+^-binding pocket (SI Fig. 32d) (21). Instead we showed – by combining NMR-experiments, site-directed point mutations and molecular dynamics simulations – that H294 is at the center of an allosteric network that contains the Ca^2+^-binding site. More specifically, in its unprotonated form H294 forms a hydrogen bond with lysine K257, which is also present in the known crystal structures of langerin (17). This hydrogen bond cannot be formed if H294 is protonated, and the allosteric mechanism that regulates the Ca^2+^-affinity likely hinges on this hydrogen bond.

Yet, protonation of H294 is only the initial detection that the surrounding medium has changed. Even though we identified the residues that are involved in the allosteric network, we do not yet understand how the protonation of H294 could ultimately affect the Ca^2+^-binding pocket. Several allosteric effects have been reported for C-type lectins (see ref. (34) for a recent review), but little is known about their underlying molecular mechanisms that could be applied to the situation in langerin. The goal of this study is to elucidate how the protonation of H294 changes the conformational ensemble of langerin, and to investigate the effect these conformational changes have on the Ca^2+^- binding pocket. A model of how the signal, that the pH has changed, traverses the allosteric network to the buried Ca^2+^-binding site and triggers the Ca^2+^-release might serve as a blueprint for understanding how pH-sensitive ligand binding is achieved in C-type lectins and other proteins.

## Results and discussion

### Structure of the langerin carbohydrate recognition domain

Langerin forms a homotrimer. The monomers consist of a short cytoplasmic tail, a transmembrane region and a long alpha-helical neck (residues 56 to 197) extending into the extracellular milieu, which carries the C-terminal carbohydrate recognition domain (17, 19). The carbohydrate recognition domain has the typical C-type lectin domain fold (Fig. 1b) (4) which consists of two extended β-sheets (turquoise), each composed of three single strands. The two β-sheets are flanked by three α-helices (purple, α_3_ in red). The carbohydrate binding pocket which contains the Ca^2+^-binding site is located on top of the β_4_-strand. One residue from this β-sheet directly binds to the Ca^2+^-ion: D308. Additionally, the Ca^2+^-ion is held in place by E293 and E285 in the long-loop (blue), which coordinate to Ca^2+^ from the side. E285 is part of a conserved EPN-motif (E285, P286, N287 in langerin), which determines the selectivity for mannose, fucose and glucose over galactose (19, 35, 36). The pH sensor H294 (pink) is located at the end of the long-loop. If its side-chain is unprotonated, it forms a hydrogen bond to K257 (also pink) in the short-loop (orange). The allosteric network that regulates the Ca^2+^- affinity comprises the long- and the short-loop (18). H229 (yellow) is the only other histidine residue in the langerin carbohydrate recognition domain. A pathogen would bind via a carbohydrate ligand (dark red) to langerin, and would be separated from the pH sensor by the long-loop. If Ca^2+^ is bound to langerin, we will call the system holo-langerin, otherwise apo-langerin.

### The effect of H294 protonation on the conformational ensemble

We conducted 31 µs of molecular dynamics simulations of holo-langerin, in which all residues were protonated according to their default protonation state at pH 7, i.e. H294 was unprotonated, and the overall protein was neutral (neutral state). We compare these simulations to 27 µs of holo-langerin, in which H294 was protonated (protonated state). Protonation of H294 has no influence on the secondary structure of langerin (Fig. 2a, Fig. 1 in the SI). Thus, any conformational change due to the protonation of H294 affects the side-chains, or those residues that are not assigned to a specific secondary structure, i.e. the loop regions.

**Figure 2.**
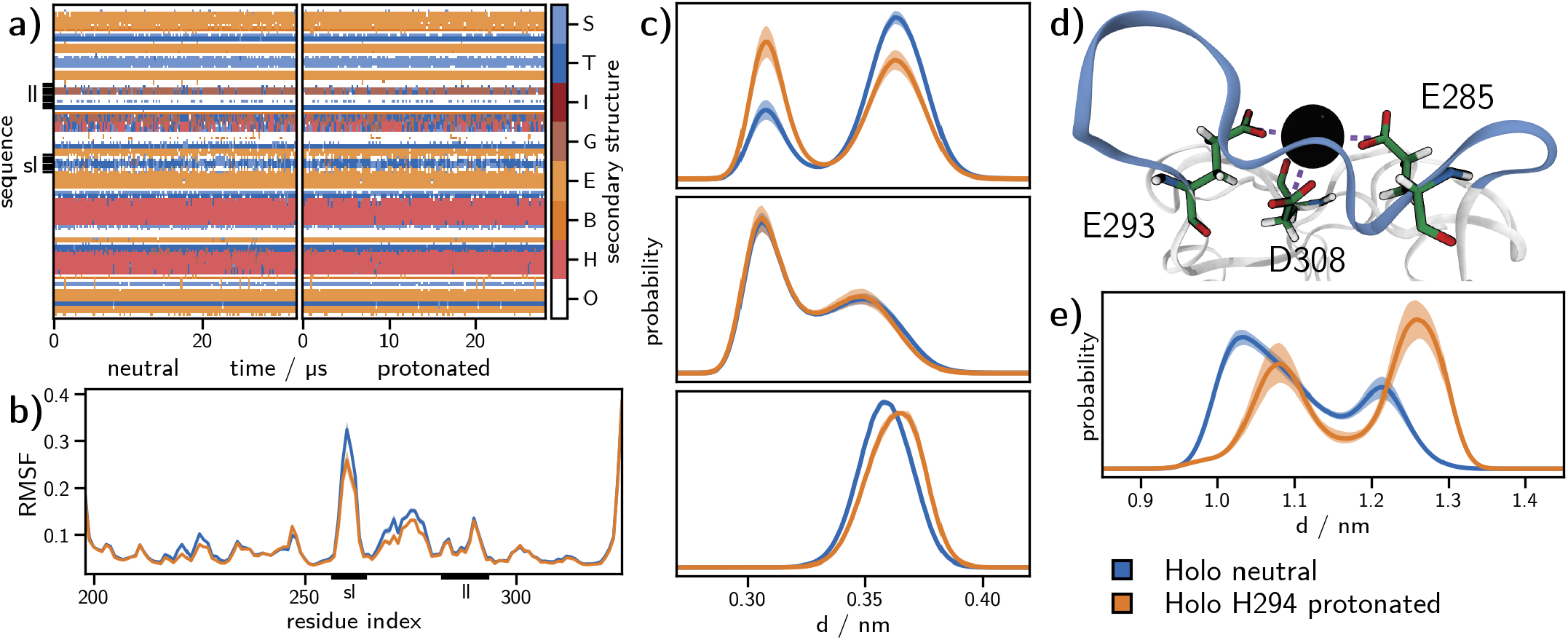
**a)** Analysis by the hydrogen bond estimation algorithm DSSP of the secondary structure in the neutral (left) and the protonated holo-state (right). Legend: S – bend, T – hydrogen bonded turn, I – 5-helix, G – 3-helix, E – extended strand, part of β-ladder, B: isolated β-bridge, H: α-helix, O: unassigned. **b)** C_α_-root-mean-square fluctuation (RMSF); sl: short-loop, ll: long-loop. **c)** Carboxyl carbon–Ca^2+^ distance histograms for E285 (upper graph), E293 (middle graph), and D308 (lower graph). **d)** Structure of the Ca^2+^-binding site showing the distances plotted in **c)** with dashed lines. **e)** Histogram of the minimum distance between Ca^2+^ and the side-chain N-atoms (N_δ_ and N_ϵ_) of H294. Solid lines: Mean of the histograms calculated for each simulation replica. Shaded area: 95 % confidence interval of the mean obtained by bootstrapping (1000 samples).

One way a conformational change in the loop regions could manifest itself, is by a change of the loop flexibility. This is however not corroborated by the root-mean-square fluctuations of the individual residues (Fig. 2b). The short-loop (sharp peak around residue 260) and the α_3_-helix (broad peak around residue 275) are more rigid in the protonated state, but the difference is very small. The flexibility in all other regions of the protein, and in particular the long-loop region, does not change upon protonation.

To gauge whether protonation of H294 has an influence on the conformation of the Ca^2+^-binding site, we measured the distance distribution between the carboxyl group of the Ca^2+^-coordinating residues – E285, E293, and D308 – and the Ca^2+^-ion (Fig. 2c–d). For E293 and D308 the differences are too minor to explain the observed difference in Ca^2+^-affinity. For E285 the distribution shifts slightly to lower distances and thus to a potentially tightly bound Ca^2+^-ion, not explaining it either. The distance difference between the two populated states is about 0.05 nm.

Yet, we know from our previous analyses (18) that protonation of H294 causes a significant shift in the conformational ensemble, and this is again confirmed by the distance distributions between the H294 side-chain and the Ca^2+^-ion in the neutral and the protonated state (Fig. 2e). In the protonated state the distribution shifts to larger distances, well beyond 1 nm. At this distance, we do not expect a significant influence of the positively charged H294 side-chain on the Ca^2+^-ion, considering that H294 is located on the protein surface and that the dielectric constant between the two interacting groups is relatively high (see SI Fig. 20 and 21 for an assessment of the Coulomb interaction) (37). Thus, we can rule out that the decrease in Ca^2+^-affinity is caused by direct Coulomb repulsion between the protonated H294 and the Ca^2+^-cation.

To uncover which residues besides H294 are involved in the conformational shift, one needs to compare the two conformational ensembles. This cannot be accomplished in the full high-dimensional conformational space. Instead one needs to project the two ensembles into a low-dimensional space that is representative of both systems. Principal component analysis (38) identifies low-dimensional spaces which preserve the directions of the largest conformational variance (39). To be able to directly compare the neutral and the protonated ensemble, we combined the simulations in the two protonation states to obtain a joint principal component space. The principal component with the largest variance represents the opening and closing of the gap between short-loop and long-loop (blue sequence of structures in Fig. 3a). The second principal component represents a sideways shear motion of the short-loop (orange sequence of structures in Fig. 3a). This is in line with our previous finding that the allosteric network is centered on these two loops (18). Even though the two principal components cover only about 28 % of the total structural variance (Fig. 3b), they represent the conformational fluctuations that are most sensitive to a protonation of H294. Separate principal component analyses of the two protonation states yielded principal components that were almost identical, indicating that the joint principal components are a faithful representation of largest variances for both protonation states. Fig. 3c shows the free energy surface of the two systems in the space of the first two joint principal components. The free-energy surface of the unprotonated system is shallow with two minima corresponding to the open and closed state of the short and the long-loop. Upon protonation, the free energy surface becomes much steeper and more structured. One can discern at least three minima. The difference plot of the probability densities in the neutral and protonated state (Fig. 3c to the right) shows these emerging conformations in red.

**Figure 3.**
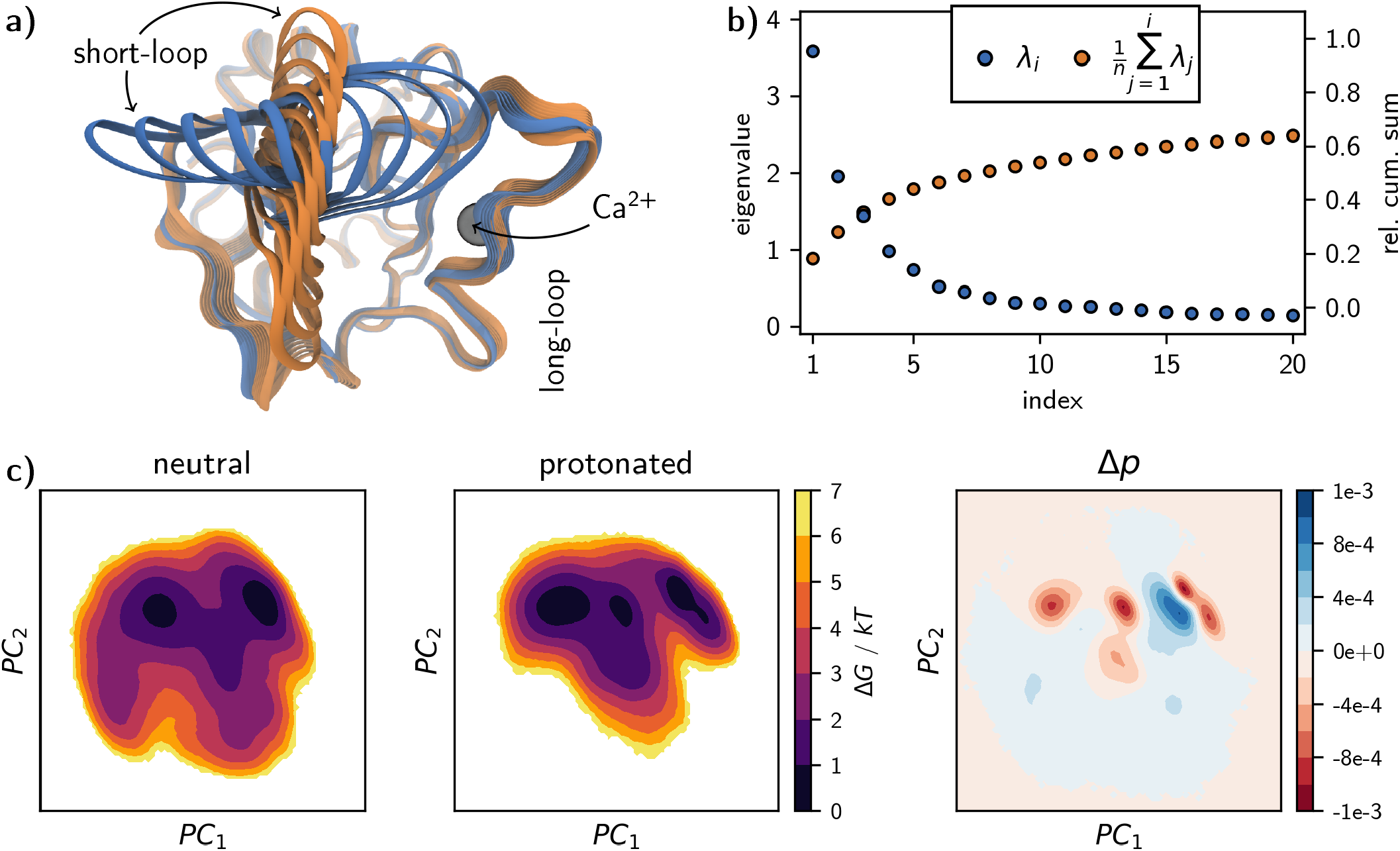
**a)** Structural interpolations along the first two principle components. **b)** Eigenvalue spectrum of the principle component analysis (blue dots) and the cumulative sum normalized by the total sum of all *N* eigenvalues 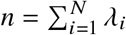 (orange dots). **c)** Free energy surfaces from the 2D projections of the individual holo-langerin trajectories onto principle components 1 and 2 and difference plot of the underlying probability distributions (neutral − protonated).

We extracted the highly populated regions by clustering in the space of the first two principal components using the density-based common-nearest-neighbors cluster algorithm (40–42), and characterized the hydrogen bond pattern of the short- and long-loop residues in each of the clusters (Fig. 4). Figs. 4c and 4d show a subset of the full analysis (see SI Fig 10) focusing on fluctuating hydrogen bonds. In the neutral state, the clusters have essentially the same hydrogen bond populations as the total ensemble, which is consistent with the shallow free-energy surface in Fig. 3c.

**Figure 4.**
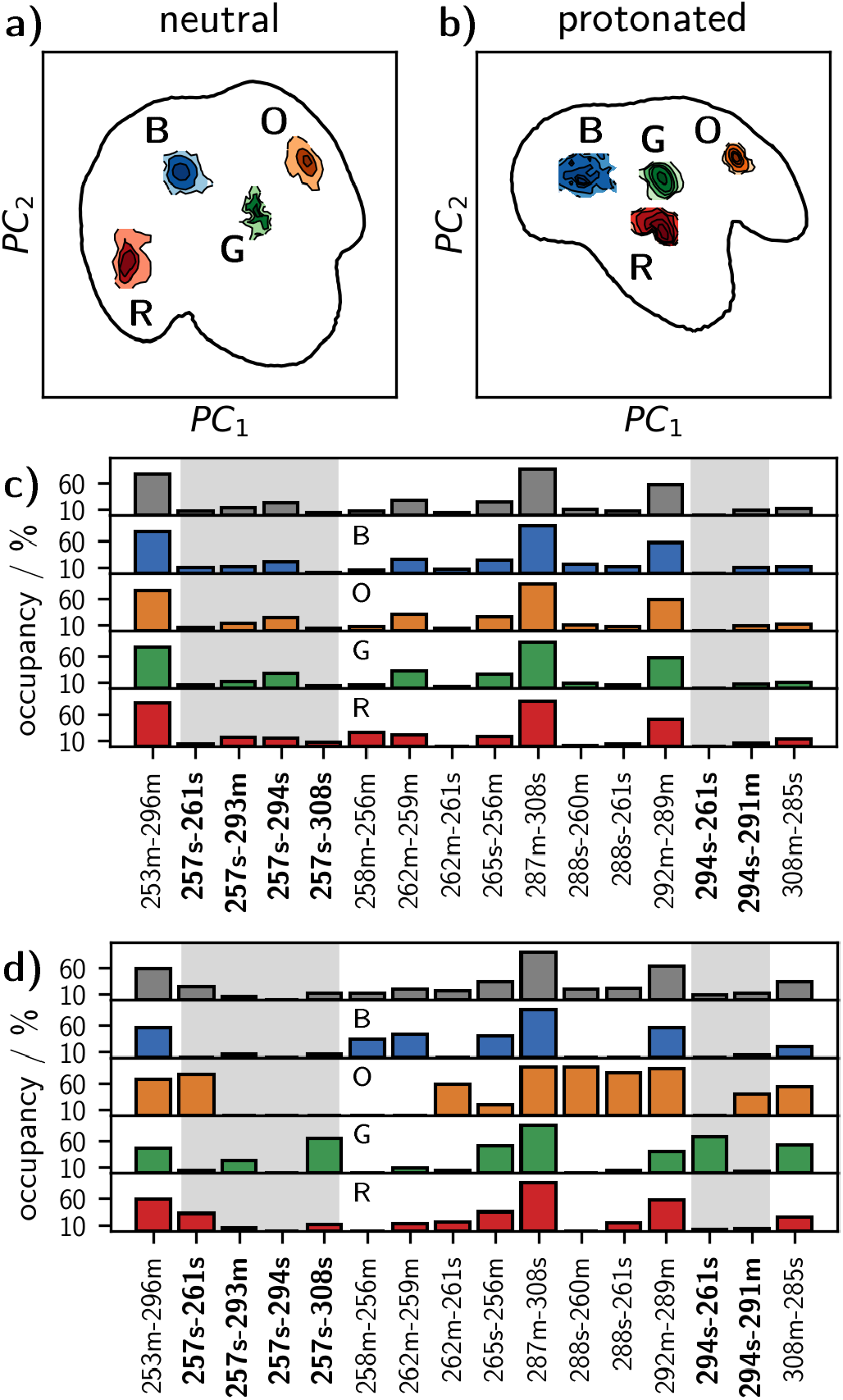
Four most populated clusters in the principal-component free energy surface of **a)** the neutral and **b)** the protonated holo-langerin. Per cluster hydrogen bond occupancy in **c)** neutral and **d)** protonated holo-langerin (populations in the full ensemble in gray). Hydrogen bonds involving K257 or H294 are highlighted by a gray background.

The situation is different in the protonated state. Here, each of the four clusters is stabilized by a hydrogen bond pattern that is distinctively different from the hydrogen bond pattern of the total distribution (Fig. 4b). This indicates that upon protonation of H294 several distinct short-loop/long-loop conformations emerge.

The most striking change arises in the green (G) cluster: the hydrogen bond between the side-chain of K257 and the side-chain of D308, which is barely populated in the unprotonated state (4.2 %), is populated to 65.4 % in this cluster (12.9 % in the ensemble). In parallel, the side-chain of the now protonated H294 forms a hydrogen bond with the carboxyl group of E261. The structure is further stabilized by a hydrogen bond between the side-chain of S265 and the main-chain of T256. Note the significance of this finding: the K257 side-chain, which is no longer engaged in a hydrogen-bond with H294, forms a new hydrogen bond with the Ca^2+^-coordinating residue D308, and thereby moves a proton into the vicinity of the Ca^2+^-binding pocket.

The conformation of the orange (O) cluster is complementary to that of the green cluster. The side-chain of K257 forms a hydrogen bond with the carboxyl group of E261, while H294 engages in a hydrogen bond to the backbone carbonyl oxygen of N291. The conformation is stabilized by hydrogen bonds between the side-chain of N288 and the backbone carbonyl oxygen of M260 and the side-chain of E261. N288 is located in the center of the long-loop, and E261 is located in the center of the short-loop. Thus, these two hydrogen bonds closely connect the two loops explaining why this structure appears in the closed-loop region of the free-energy surface. The main-chain-main-chain hydrogen bond between N292 and A289 additionally stabilizes this structure.

The blue (B) cluster is an open-loop structure in which neither K257 nor H294 are engaged in one of the considered hydrogen bonds. It features the 258m–256m and 262m– 259m hydrogen bonds within the short-loop. The red (R) cluster is a slightly sheared structure in which the K257 side-chain partly forms a hydrogen bond to the carboxyl group of E261 and partly to the carboxyl group of D308.

Three hydrogen bonds in Fig. 4 directly involve Ca^2+^- coordinating residues. First, we already discussed the hydrogen bond K257–D308. Second, the hydrogen bond between the main-chain of N287 and the side-chain of D308 is important for the stability of the long-loop fold. It is occupied to about 90 % in both protonation states. Third, population of the hydrogen bond between the main-chain amid group of D308 and the carboxyl group of E285 is increased in the protonated state. This is particularly true for cluster G (green) and O (orange). This hydrogen bond might compete with the coordination of E285 to Ca^2+^ and thereby might contribute to the observed decrease in Ca^2+^- affinity. In both, the neutral and the protonated system, the bonds N288s–M260m, N288s–E261s, K257s–E261s and G262m–E261s are strongly correlated (see SI Fig 10). In the protonated state a strong correlation between K257s–D308s and H294s–E261s arises, indicating that these two hydrogen bonds are formed and broken simultaneously.

### A mechanism for the pH-sensitive Ca^2+^-affinity in langerin

We are now ready to propose a mechanism that explains how protonation of H294 can lead to a decrease in Ca^2+^-affinity. In the neutral state, K257 and H294 form a hydrogen bond which is populated over a wide range of conformations. We also observe a weak hydrogen bond of the K257 side-chain to the main-chain of the Ca^2+^-coordinating residue E293, but direct hydrogen bonds to the Ca^2+^-coordinating carboxyl groups are hardly ever formed (Fig. 5a). Upon a drop of pH from 7 to 6, the side-chain of H294 is protonated in accordance with its p*K*_a_: H294 is the initial pH sensor. The protonation of H294 changes the hydrogen bond pattern between the short and the long-loop. In particular the side-chains of H294 and K257 form new contacts, which gives rise to previously inaccessible conformations. Cluster O (orange) and cluster G (green) exhibit mutually exclusive hydrogen bond patterns. In cluster O, multiple hydrogen bonds connect the short and the long-loop causing a closed loop conformation. The positively charged side-chain of K257 forms a hydrogen bond to the negatively charged side-chain of E261. But similar to the neutral state, there is no direct hydrogen bond to the Ca^2+^-coordinating carboxyl groups (Fig. 5c). This is different in cluster G. Here the positively charged side-chain of H294 forms a hydrogen bond with the negatively charged carboxyl group of E261. Simultaneously, the positively charged side-chain of K257 forms a hydrogen bond with the carboxyl group of D308 (Fig. 5b). This hydrogen bond withdraws electron density from the coordinative bond between D308 and Ca^2+^, and thereby reduces the Ca^2+^-affinity. It is even conceivable that the proton is transferred entirely to the carboxyl group of D308 (43). We thus propose that cluster G (green) is responsible for the decrease in Ca^2+^-affinity at pH 6.

**Figure 5.**
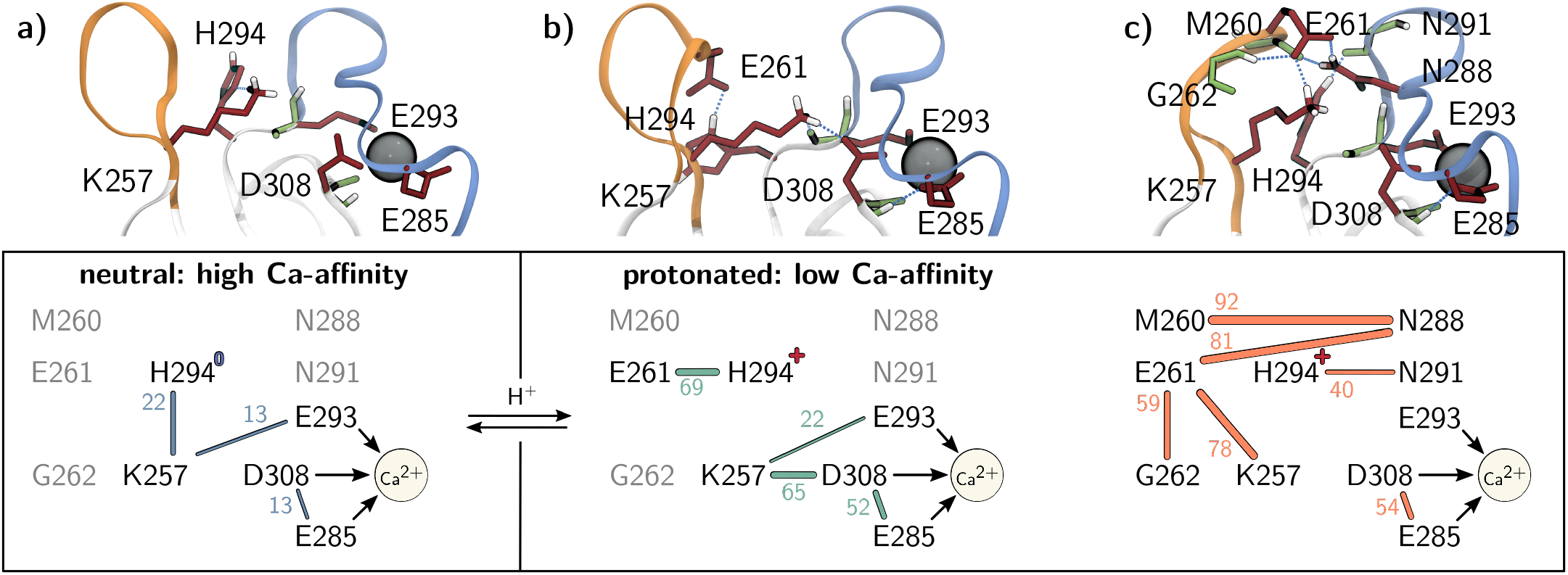
Allosteric mechanism for the pH-sensitive Ca^2+^-affinity in langerin. **a)** neutral state, **b)** cluster G (green) in the protonated state, and **c)** cluster O (orange) in the protonated state. Lines: hydrogen bonds with population in percent. Arrows: Coordination between carboxyl groups and Ca^2+^.

In this mechanism, K257 acts as a proton reservoir. The initial detection of a pH change via protonation of H294 leads to the cluster G, in which K257 moves a proton into the vicinity of the Ca^2+^-binding site and locally increases the proton concentration. Thus, the signal that the pH has changed is allosterically transferred to the Ca^2+^-binding pocket without transferring the actual proton that triggered the mechanism.

A crucial assertion in the proposed mechanism is, that the life-time of cluster G (green) represents a distinct conformation, that is stable enough for the Ca^2+^-ion to leave the binding pocket. The fact that cluster G corresponds to a free-energy minimum in the space of the principal components hints at a stable conformation. But because the principal components maximize the spatial variance and not the variance in time, this is not sufficient to be certain.

Fig. 6a shows the distance distribution between the K257 and D308 side-chain for the neutral and the protonated state. In both protonation states, the maximum at short distances around 0.2 nm is well separated from the maximum at larger distances.

**Figure 6.**
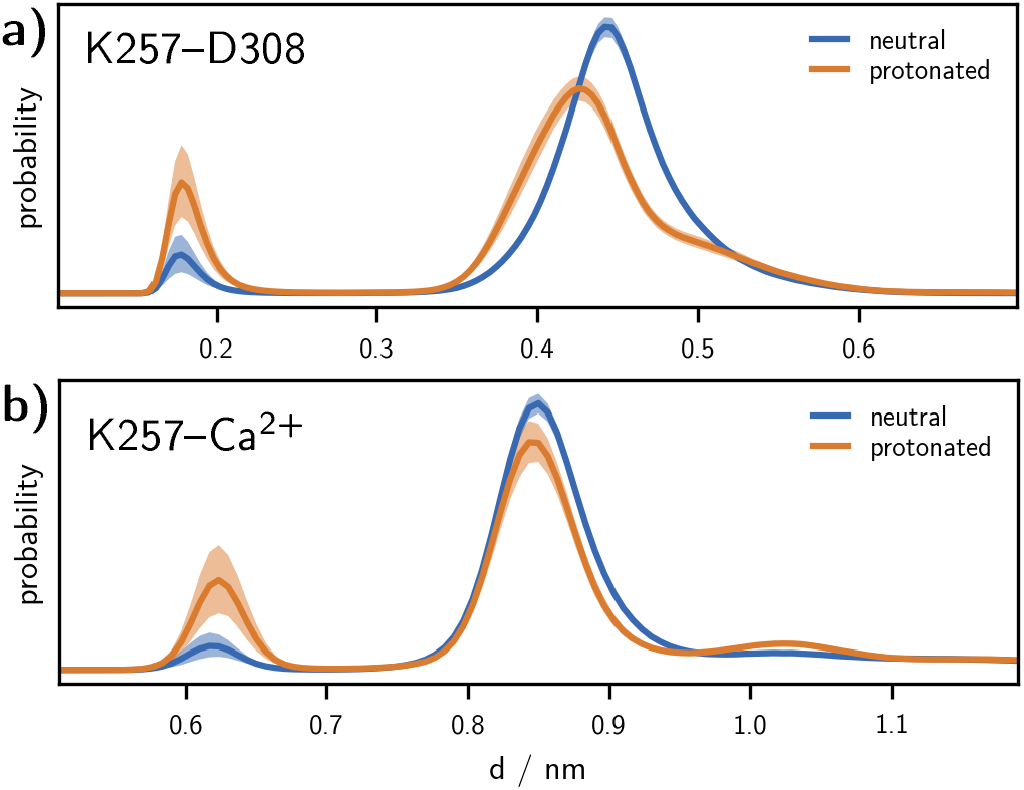
**a)** K257–D308 side-chain distance distribution. **b)** K257 side-chain amine – Ca^2+^distance distribution. Solid lines: Mean of the histograms calculated for each simulation replica. Shaded area: 95 % confidence interval of the mean obtained by bootstrapping (1000 samples).

In the neutral state, the short distances are populated only in 4.3 % of all simulated conformations, which increases to 13.2 % when H294 is protonated. This is in line with the increase of population in the K257–D308 hydrogen bond from 4.2 % to 12.9 %. We obtain the same results, when plotting the distance between the K257 side-chain amine and the Ca^2+^-ion in Fig. 6b. Thus, cluster G (green) indeed represents a distinct conformation.

To assess the stability of conformations in cluster G (green), and to relate its formation to other dynamic processes in the protein, we constructed a core-set Markov model of the conformational dynamics (44–46). In Markov models, the conformational space is discretized into states and the conformational dynamics are modeled as Markov transitions within a lag time *τ* between pairs of these states, where the transition probabilities are obtained from molecular dynamics simulations. From the eigenvectors and eigenvalues of the Markov-model transition matrix one obtains long-lived conformations as well as the hierarchy of the free-energy barriers separating them. The special feature of core-set Markov models is, that the states are confined to the regions close to the minima of the free-energy surface, i.e. so-called core-sets, whereas the regions between these minima are modeled by committor functions. This reduces the discretization error of the model considerably.

The Markov-model construction is preceded by a dimensionality reduction of the conformational space using the time-independent component analysis (47, 48). Time-independent components (tIC) maximize the variance within lag-time *τ* rather than the instantaneous variance maximized by principal components. A projection into a low-dimensional tIC-space can thus be interpreted as projection into the space of the slowly varying coordinates of the system. Fig. 7a shows the free-energy surface of the protonated system projected into the space of the first and the second tIC (see SI for other projections), and Fig. 7b shows the projection of cluster G (green) and O (orange) into this space.

**Figure 7.**
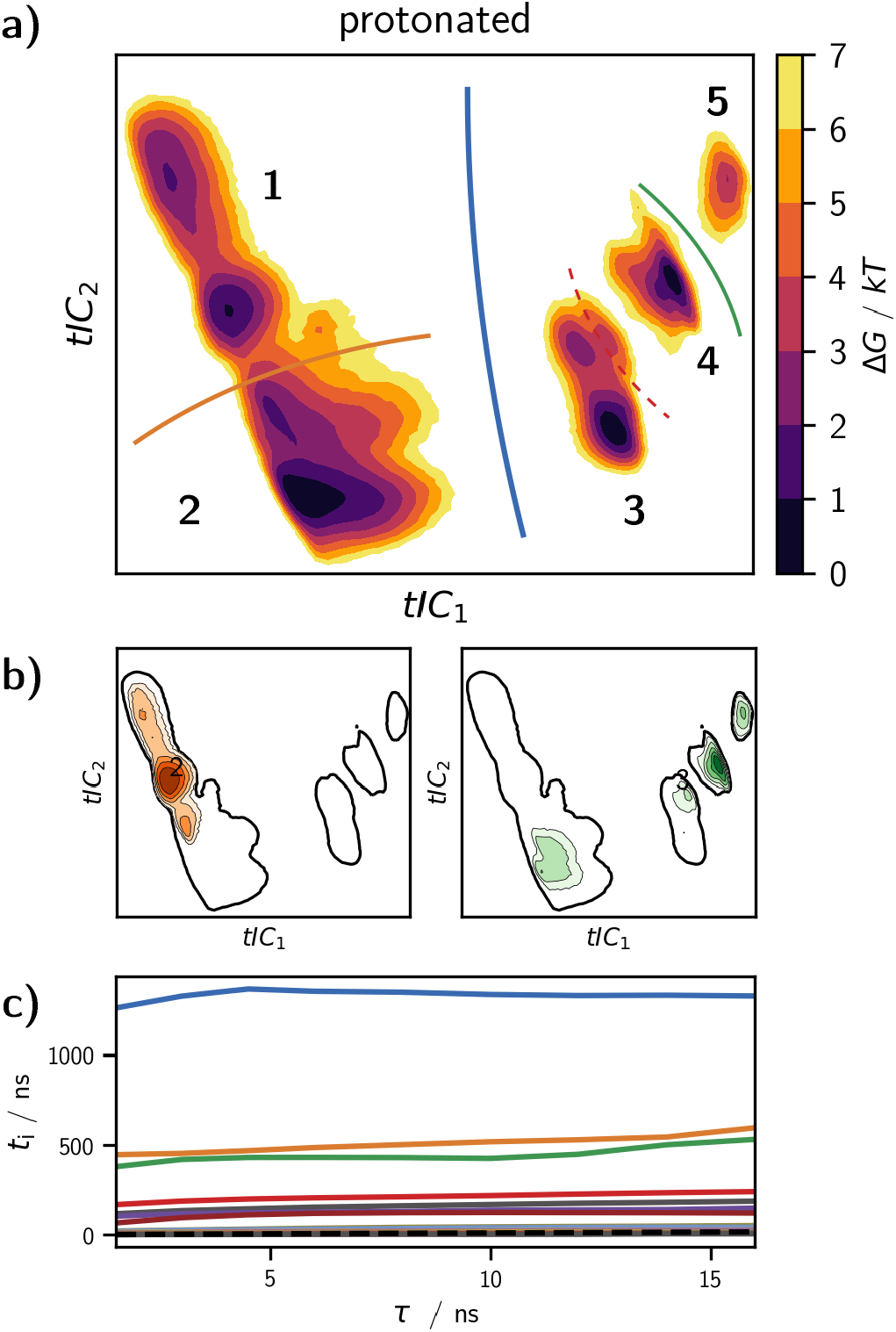
Core-set Markov model of the conformational dynamics of protonated holo-langerin. **a)** Free energy surfaces from the 2D projections of protonated holo-langerin trajectories onto the first two tICs. Solid lines: transition regions between the five meta-stable states connected by the four slowest dynamic processes. **b)** Projections of cluster G (green) and O (orange) into the space of the first two tICs. **c)** Implied time scales of the core-set Markov model. The colors of the processes match the transition regions drawn into **a)**.

We then identified 22 core-sets in the space of the first six tICs using common-nearest-neighbors clustering (40–42), and used them to construct a core-set Markov model. The implied time-scale test shows that the timescales of our core-set Markov model are independent of the lag time *τ* indicating a very small discretization error and thus a high-quality Markov model (Fig. 7c). The slowest dynamic process occurs on a timescale of about 1.3 µs and corresponds to changes in the local conformations of E261 and its hydrogen bond pattern. It thus separates the conformations of cluster G (green) and cluster O (orange) along the blue barrier in Fig. 7a. Note that all conformations in which the K257s–D308s hydrogen bond is formed alongside H294s–E261s are located on the right-hand side of this barrier (see SI). The fact that we find some structures that have originally been assigned to the G (green) conformation on the left-hand side of the barrier is likely due to the insufficient separation of long-lived conformations in the principal component space (Fig. 7b). Next, protonated langerin has two slow timescales that occur at about 500 ns. One process describes transitions between the closed-loop conformations in region 1 and conformations in which the distance between the long and the short-loop is larger in region 2. The other process represents a transition between conformations in which the backbone-orientation of N291 forbids the N292m–A289m hydrogen bond giving rise to a distortion of the long-loop (region 5) and the conformations in which the N292m–A289m hydrogen bond is possible (regions 3 and 4). The dashed barrier marks transitions to more open short-loop forms occurring on a timescale of 210 ns.

In summary, conformations in which the K257–D308 hydrogen bond is formed are separated from the alternative O (orange) conformation by a rare transition that occurs on a timescale of 1.3 µs. Within the right-hand side of the barrier in Fig. 7a the G (green) conformation is at least stable on a timescale of 200 ns. This is likely sufficient to enable the escape of the Ca^2+^-ion from the binding pocket. A core-set Markov model of neutral holo-langerin is reported in the SI.

To directly probe how the stability of the Ca^2+^-bound state of the protein depends on the protonation state and on the conformation of langerin, we used constant-velocity steered-MD experiments (49–51). In these simulations, a force that increases linearly with time is applied to the Ca^2+^-atom (Fig. 8a), and the opposing force (i.e. the resistance against this pulling force) is measured. At a certain maximum force the ionic bonds between the Ca^2+^-atom and the coordinating residues rupture and the Ca^2+^-ion leaves the binding pocket. In the computer experiment, this is marked by a sudden drop in the opposing force (Fig. 8c). The rupture force is a rough measure for the free-energy difference to the transition state Δ*G*^‡^. The rational is that a deeper free-energy minimum of the Ca^2+^-bound state is associated with a steeper slope to the transition state, and the rupture force, reflecting the maximal slope, reports on the stability of the Ca^2+^-bound state (52, 53). We chose the pulling rate such that the rupture events are observed after several nanoseconds. This ensures, that the system has enough time to adjust to the pulling, but also that the initial starting conformation is preserved to some degree.

**Figure 8.**
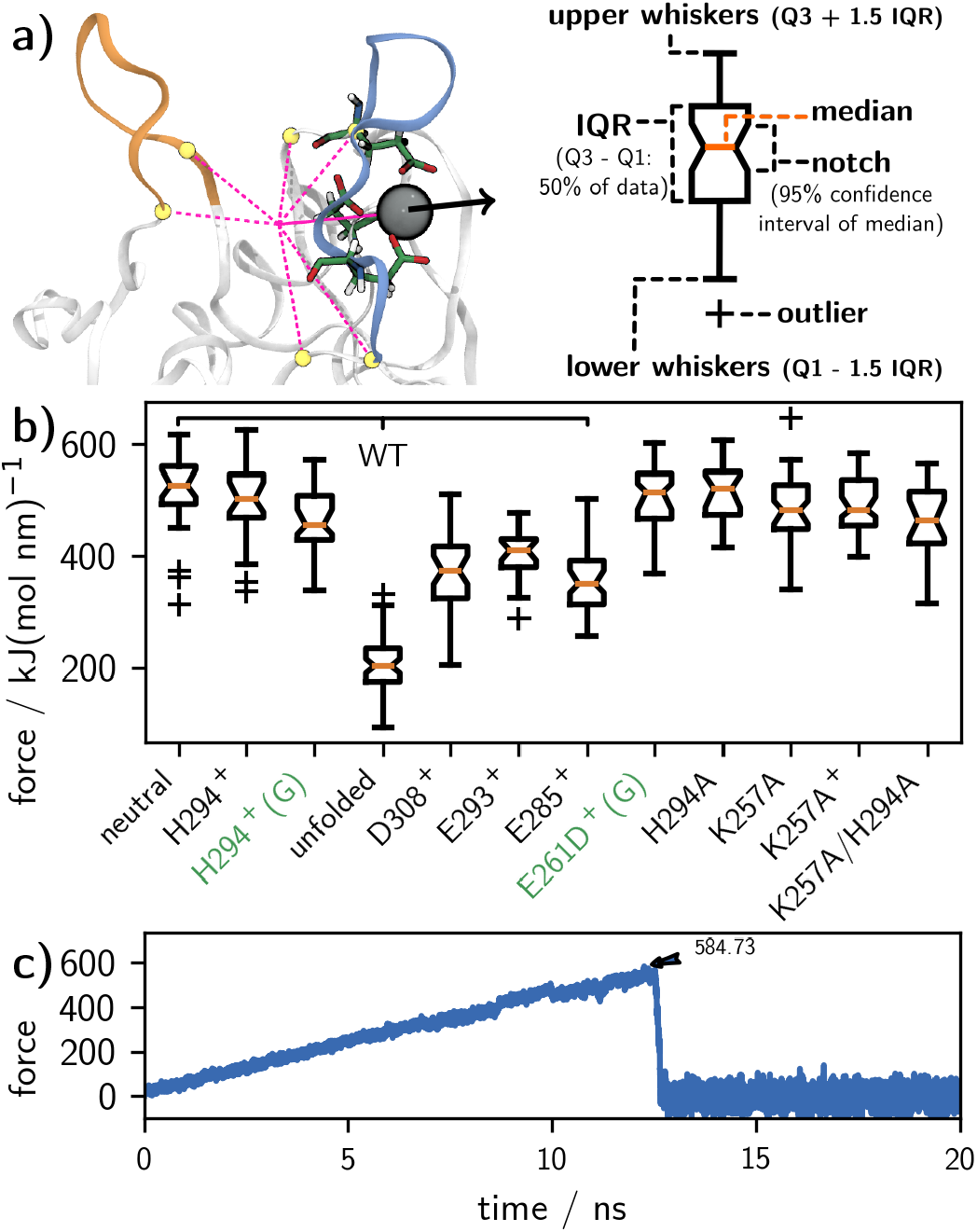
**a)** Pull coordinate defined as the distance vector between Ca^2+^ and the center-of-mass of the C_α_-atoms of residues 257, 264, 281, 282, 293, and 294. **b)** Maximal pulling force observed acting on Ca^2+^during simulations of langerin in various states as notched box representation. The orange line represents the median, while the box enframes the interquartile range. The box notches indicate the 95 % confidence interval on the median. Points lying beyond 1.5 times the edges of the box are regarded as outliers (**+**), and the whiskers mark the data range without outliers. **c)** Example for a force trajectory with a rupture event at about 12 ns. Maximum force indicated by an arrow.

For each system, we conducted 40 steered-MD simulations and report the data as notched boxplot in Fig. 8b. Overall, the plot shows that we could determine the median of rupture force with high confidence and hardly any outliers. The rupture force decreases from the neutral to the protonated system (H294^+^) and then further to simulations of the protonated system started in the G (green) conformation, in which the K257 amine forms a hydrogen bond with the D308 carboxyl group. This decrease is predicted by our mechanism. Note that classical force-fields cannot model instantaneous shifts in the electron density due to the formations of hydrogen bonds. Thus, the rupture force in the G (green) conformation might actually be somewhat lower. If the Ca^2+^-coordinating residue D308 is protonated, corresponding to a situation in which the proton is transferred from K257 to D308, the rupture force is about 150 kJ/(mol nm) lower than in the neutral system.

The same is observed when one of the other two Ca^2+^- coordinating residues is protonated. A drastic reduction in the rupture force is observed, when the experiment is started from a state where the long-loop is unfolded. This is expected, as one of the Ca^2+^-coordinating residues E285 is removed from the cage of the binding site in this arrangement. The rupture force for the mutant E261D (started from an analogon of the G conformation) and the mutant H294A are in the same range as for the neutral wild-type langerin.

Notably, the rupture forces for K257A mutants are insensitive towards the modeled state of H294. The binding capability is virtually the same, no matter if H294 is neutral, protonated or mutated. This substantiates the importance of K257 to transport a protonation signal to the Ca^2+^-binding site.

### Comparison to experimental data

The Ca^2+^-dissociation constant of wild-type langerin at pH 7 is *K*_d_ = 105 ± 15 µM, and increases to *K*_d_ = 800 ± 150 µM at pH 6 (18), as determined by ITC. These dissociation constants correspond to binding free energies of Δ*G*_pH 7_ = –22.9 kJ/mol, and Δ*G*_pH 6_ = – 17.8 kJ/mol at *T* = 300 K, yielding a pH sensitivity of ΔΔ*G* = Δ*G*_pH 6_ – Δ*G*_pH 7_ = 5.1 kJ/mol (Fig. 9). By contrast the dissociation constant of the H294A mutant, in which the pH sensor H294 is removed, are *K*_d_ = 35 ± 15 µM at pH 7 (Δ*G*_pH 7_ = – 25.6 kJ/mol), and *K*_d_ = 125 ± 5 µM at pH 6 (Δ*G*_pH 6_ = – 22.4 kJ/mol), corresponding to a reduced pH sensitivity of ΔΔ*G* = 3.2 kJ/mol (18) (Fig. 9). Our mechanism so far explains the pH sensitivity due to the pH sensor H294. The fact that the H294A mutant exhibits a residual pH-sensitivity indicates that langerin has a second pH sensor.

**Figure 9.**
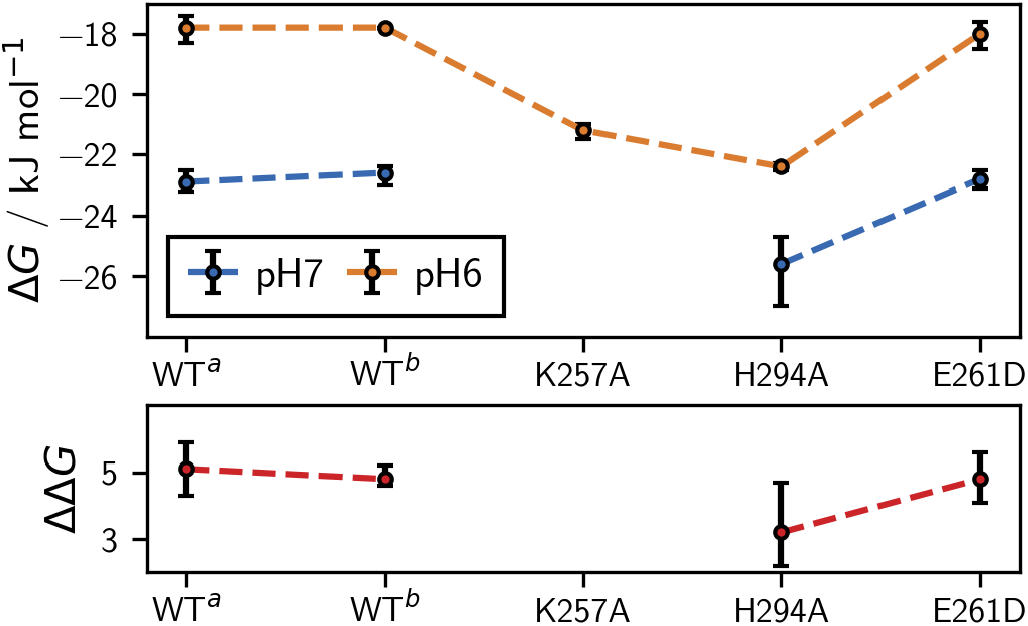
Ca^2+^-binding free energies under standard conditions in kJ/mol, calculated as Δ*G* = – *RT* ln (*K*_d_), where *R* = 8.314 J/(K mol) is the gas constant, *T* = 300 K is the temperature, and *K*_d_ in units of mol/L are the experimentally determined dissociation constants. Measurements at pH 6 (blue) and pH 7 (orange) with experimental uncertainties indicated with errorbars and pH-sensitivities in kJ/mol calculated as ΔΔ*G* = Δ*G*_pH 6_ −Δ*G*_pH 7_ (red).

To convince ourselves of the robustness of these results, we remeasured the dissociation constants of wild-type langerin (see SI). We obtained *K*_d_ = 113 ± 14 µM at pH 7 (Δ*G*_pH 7_ = – 22.6 kJ/mol), and *K*_d_ = 802 ± 150 µM at pH 6 (Δ*G*_pH 6_ = – 17.8 kJ/mol), yielding a pH sensitivity of ΔΔ*G* = 4.8 kJ/mol (Fig. 9). This is in excellent agreement with our previous results.

Four residues are central to our mechanism: H294, K257, D308, and E261. D308 directly coordinates to Ca^2+^and is therefore not a suitable candidate for site-directed mutagenesis. In contrast to H294A, the pH-sensitivity of K257A could not be determined because the protein precipitated at pH 7. However, both mutants have a higher Ca^2+^-affinity than wild-type langerin at pH 6, which previously could not be explained. The overall higher Ca^2+^-affinity in the K257A mutant is predicted by our mechanism, because the K257–D308 hydrogen bond that destabilizes the Ca^2+^- coordination cannot be formed in the absence of the K257 side-chain. The H294A mutant has the K257 side-chain, and the conformation in which K257 is in the vicinity of D308 (Fig. 6) can in principle be formed. However, in our simulations of H294A we find that the K257 side-chain is in the vicinity of the D308 side-chain in only 1.7 % of the simulated structures, which might explain the higher Ca^2+^-affinity of the H294A mutant (see SI).

Besides H294 and K257, residue E261 is important for the stabilization of the G (green) conformation, which is responsible for lowering the Ca^2+^-affinity. However, it also stabilizes the cluster O (orange), which is not expected to increase the Ca^2+^-affinity, because K257 forms a hydrogen bond with E261 rather than with D308 in this conformation. We therefore predicted that mutating E261 has little effect on the pH-sensitivity. We measured the Ca^2+^-dissociation constants for the E261D mutant at pH 6 and pH 7 by ITC (see SI), and the results confirm our prediction. The dissociation constants of the E261D mutant are *K*_d_ = 108 ± 11 µM at pH 7 (Δ*G*_pH 7_ = –22.8 kJ/mol), and *K*_d_ = 742 ± 141 µM at pH 6 (Δ*G*_pH 6_ = –18.0 kJ/mol), yielding a pH sensitivity of ΔΔ*G* = 4.8 kJ/mol (Fig. 9).

### Long-loop unfolding

So far, our mechanisms explains how Ca^2+^ is destabilized in the binding pocket of holo-langerin. However if the proton is transferred from K257 to D308, the mechanism also has profound effects on apo-langerin. In holo-langerin the long-loop is stabilized in a well-defined conformation (folded long-loop conformation) by E285 which coordinates to Ca^2+^. In apo-langerin this interaction is not possible, and the long-loop spontaneously unfolds in our simulations. Similar long-loop unfolding has been observed in the crystal structures of other C-type lectins, like tetranectin (54), TC14 (55) or MBP (56). To estimate the unfolding rate we conducted 30 to 60 simulations (see SI) for each of the following protonation states of apo-langerin: neutral, H294 protonated, H294 and E285 protonated, H294 and E293 protonated, and H294 and D308 protonated, each of them started in the folded conformation. In four of the five protonation states 44 to 54 % of all trajectories unfold within 220 ns simulation time, as determined by visual inspection (Fig. 10c, blue dots). The carboxyl group D308 is critical for the stabilization of the folded loop conformation in the absence of Ca^2+^ by forming hydrogen bonds with the N287 side-chain, as well as with the backbone amide-hydrogen of N287 and N288 (Fig. 10a). If D308 is protonated, all three hydrogen bonds are much weaker, and consequently the long-loop unfolds at a higher rate (75 % within 220 ns).

Long-loop unfolding often occurs via an intermediate conformation, in which the hydrogen bonds with the backbone amides of N287 and N288 are broken, while the hydrogen bond to the N287 side-chain is still possible. In this intermediate form the loop is more flexible than in the fully folded state, but the characteristic turns in the loop backbone are still largely present, and we observe refolding to the fully folded state in some of the trajectories. The transition to the fully unfolded conformation occurs when one or more of the backbone torsion angles in the long-loop rotate, and the hydrogen bond between the side-chains of D308 and N287 breaks. This transition is irreversible on the timescale of our simulations.

**Figure 10.**
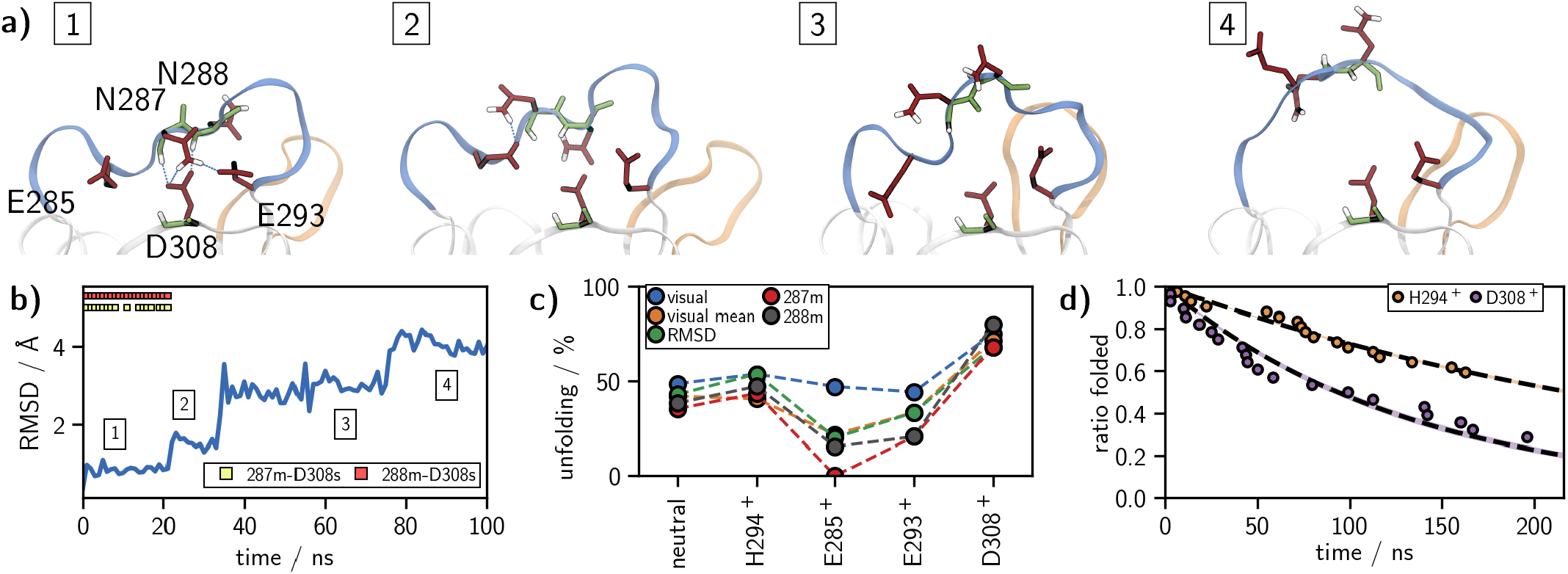
**a)** Long-loop unfolding in apo-langerin with example structures for a 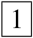 fully folded, 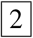 intermediate, 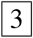 partially unfolded, and 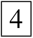 fully unfolded state. **b)** Example trajectory of the long-loop C_α_-RMSD, 22 ns: intermediate state, 30 ns: unfolding event **c)** Percentage of unfolded trajectories within 220 ns determined by: last folded frame (visual), mean of last folded and first unfolded frame (visual mean), RMSD > 0.2 nm, and hydrogen bonds N287m–D308s (287m) and N288m–D308s (288m). **d)** Decay plot of folded trajectories (last folded frame) and exponential fit (dashed line ±*σ*), H294^+^: H294 protonated, D308^+^: H294 and D308 protonated.

To corroborate our visual analysis of the simulation end points, we determined the time of the unfolding event by four additional criteria: the mean between last fully folded frame and first fully unfolded frame determined by visual inspection, the C_α_-RMSD of the long-loop residues exceeds 0.2 nm, and breaking of the hydrogen bonds between the D308 carboxyl group or the backbone amide-hydrogen of N287 and N288. All four criteria confirm the first analysis (Fig. 10c). If E285 is protonated a hydrogen bond between the protonated carboxyl group of E285 and the unprotonated carboxyl group of D308 stabilizes a partially folded loop structure, such that for some criteria we observe even fewer unfolding events than by the simple visual analysis for this system. We determined the half-life periods *t*_1/2_ of the folded states from the decay plots of the folded trajectories (see SI). Independent of the criterion, the decay is fastest, when D308 is protonated. In particular unfolding is over twice as fast if D308 is protonated than if only H294 is protonated (*t*_1/2_ = 218 versus 93 ns, Fig. 10d). Some of the decays deviate from a single-exponential decay, hinting at a more complex underlying unfolding mechanism.

Since the folded conformation binds Ca^2+^ much more strongly than the unfolded conformation (Fig. 8), the equilibrium between folded and unfolded long-loop is critical for the overall Ca^2+^-affinity. Thus, the protonation of D308 has a two-fold effect: First, it destabilizes the Ca^2+^-ion in the binding pocket. Second, after the Ca^2+^-ion has left the binding pocket, it destabilizes the folded loop conformation and thereby reduces the likelihood of Ca^2+^-rebinding.

### The second pH sensor

In the ITC experiments the H294A mutant exhibits a pH sensitivity of Δ*G* = 3.2 kJ/mol even though the pH sensor H294 is missing (18). This suggests that langerin has a second pH sensor. To convince ourselves that this residual pH sensitivity is indeed due to a second pH sensor, we checked whether K257 forms another potentially pH-sensitive hydrogen bond in the H294A mutant which could replace the pH-sensitive K257–H294 hydrogen bond and explain the residual pH sensitivity. In our simulations of the H294A mutant, K257 does not form any highly populated hydrogen bond. With 13 % population the hydrogen bond between the side-chain of K257 and the main-chain carbonyl group of E293 is the most frequently formed hydrogen bond. However, in wild-type langerin it is formed with the same frequency. All other hydrogen bonds of K257 are populated with less than 5 %. Thus, the experimentally determined pH-sensitivity in the H294A mutant does indeed indicate that wild-type langerin has a second pH sensor.

There are two possible mechanisms to explain the residual pH-sensitivity. First, langerin could have a second allosteric pH sensor that, similar to H294, is activated by protonation from the surrounding solvent prior to the dissociation of Ca^2+^. Second, the carboxyl groups of the Ca^2+^-coordinating residues E285, E293, and D308 could form a dyad with an effective p*K*_a_ that makes it sensitive to a pH change form 7 to 6. That is, after initial dissociation of Ca^2+^, one of the coordinating residues (Fig. 2d) is protonated and the protonated state is stabilized as a hydrogen bond to an unprotonated carboxyl group (57). We first discuss the possibility of a second allosteric pH sensor before investigating whether a dyad is possible.

H229 is the only other histidine residue in langerin. It is solvent exposed and will indeed be protonated when the pH changes from 7 to 6. However, H229 is located far away from the Ca^2+^-binding site which makes an allosteric influence on the Ca^2+^-binding affinity unlikely (Fig. 1). This is further corroborated by the previously published mutual information analysis of the allosteric network in langerin and by chemical shift perturbation experiments (18). In the extended simulation data set used for this study, protonation of H229 has a local effect on the “lower” protein region including the α_1_-helix, but these conformational shifts are well separated from the Ca^2+^-binding site. We therefore exclude H229 as a potential pH-sensor.

Other candidates for allosteric pH sensors are aspartic and glutamic acids, whose p*K*_a_ (in water at 25 °C 4.15 for E and 3.71 for D) (28) can be shifted by several p*K*_a_ units by the local environment in the protein, such that their carboxyl groups could become sensitive to a pH change from 7 to 6 (58). Apart from the Ca^2+^-coordinating residues E285, E293, and D308, langerin has nine aspartic or glutamic acids. Using PROPKA 3.1 (59, 60), we calculated the distribution of the p*K*_a_-values for these residues in holo-langerin in the neutral and the H294-protonated state, as well as for apo-langerin in the neutral and the H294-protonated state. The distributions are based on 10,000 to 30,000 structures extracted from the simulations of the corresponding systems, and are reported along with the mean and the standard deviation in the SI. The mean p*K*_a_-value of all tested residues is below 5.0, and none of the distributions reaches into the critical region between pH 6 and 7, indicating that none of them acts as pH sensor. We therefore conclude that the residual pH-sensitivity in langerin is not generated by a second allosteric pH-sensor.

PROPKA 3.1 can detect the coupling between two carboxyl groups that are in close vicinity. It returns two alternative p*K*_a_-values. In alternative *a*, one carboxyl group is protonated first and stabilized by the second (unprotonated) carboxyl group, in alternative *b* the situation is reversed. Fig. 11a shows the p*K*_a_-distribution of the Ca^2+^- coordinating residues E285, E293, and D308 as well as the p*K*_a_-distribution of H294 for apo-langerin in the neutral and the H294-protonated state. No coupling between E285, E293, and D308 was detected by PROPKA 3.1. Their mean p*K*_a_-value is below 5.0, and none of the distributions reaches into the critical region between pH 6 and 7. By contrast, the mean p*K*_a_-values of H294 are about 6 in the neutral and the H294-protonated state, and the p*K*_a_-distributions have a large overlap with the critical region between pH 6 and 7. Thus, from these simulations one would conclude, that langerin does not have a protonatable dyad in the Ca^2+^-binding pocket, and that only H294 is sensitive to a pH change from 7 to 6.

**Figure 11.**
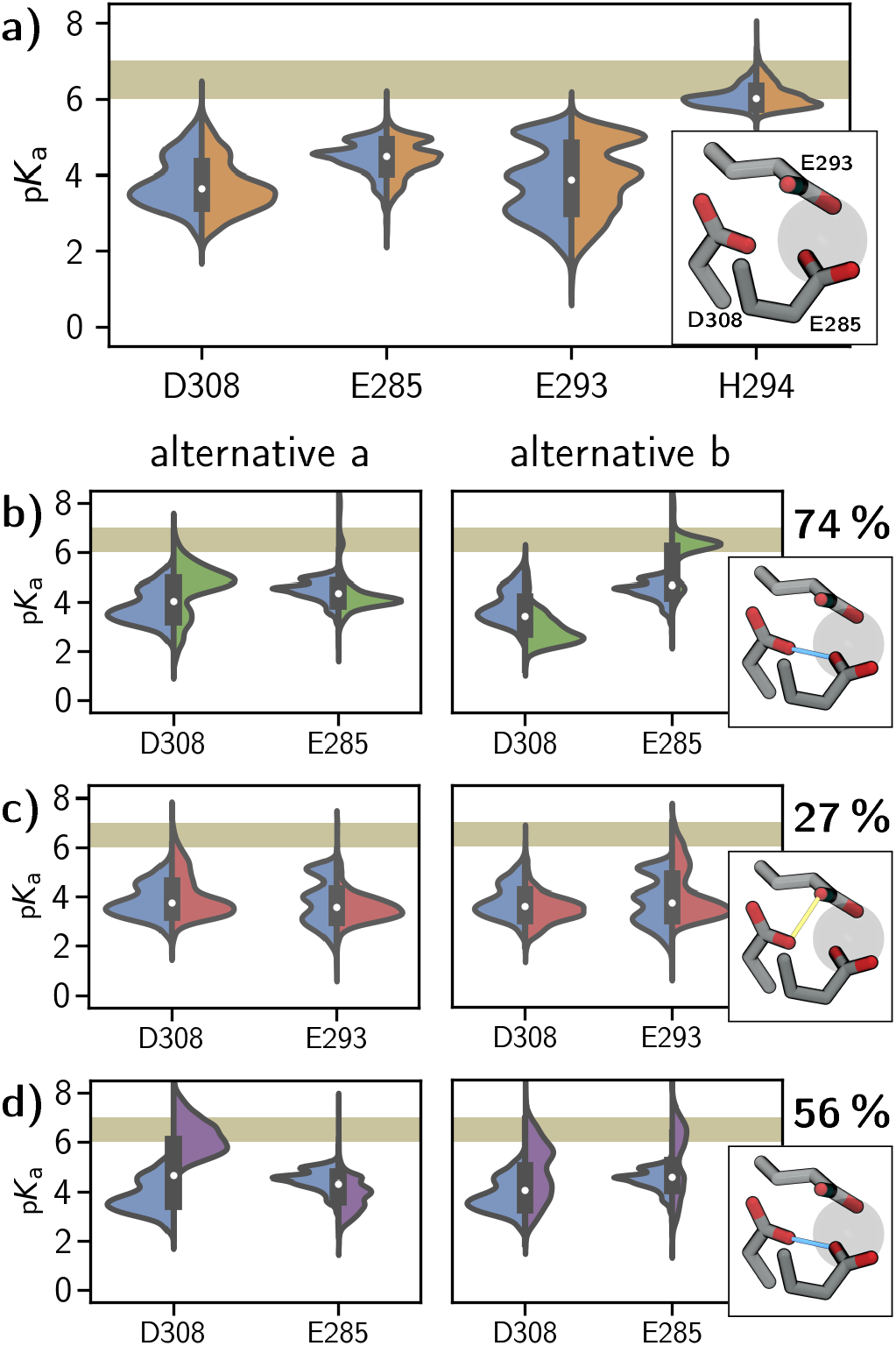
p*K*_a_-Value distributions (all frames considered) calculated with PROPKA 3.1 for **a)** the neutral apo- (blue) and the H294 protonated (orange) system. Distributions for residues involved in coupling in **b)** neutral versus E285 protonated (green), **c)** neutral versus E293 protonated (red), and **d)** neutral versus D308 protonated (purple). Alternative distributions due to the coupling left and right. Percentages of coupling frames are placed over the binding site illustrations.

However, in the neutral and the H294-protonated state, the carboxyl group of the Ca^2+^-coordinating residues are negatively charged and repel each other, making structures in which the two carboxyl groups are close enough to potentially stabilize a protonation unlikely. We therefore also calculated the p*K*_a_-distribution for the following protonation states of apo-langerin: H294 and E285 protonated (Fig. 11b), H294 and E293 protonated (Fig. 11c), and H294 and D308 protonated (Fig. 11d). For these protonation states, substantial coupling between the Ca^2+^-coordinating residues is detected. D308 and E285 couple in 74 % of all structure if E285 is protonated, and in 56 % of all structures if D308 is protonated. When E293 is protonated, E293 and D308 couple in 27 % of all structures.

These couplings give rise to a strong shift of the p*K*_a_ distributions compared to neutral apo-langerin. We report the distributions of both p*K*_a_-estimates, which should be interpreted as limiting cases of the true distribution. If D308 is protonated, the p*K*_a_-distributions of D308 for both limiting cases reach well into the critical region between pH 6 and 7, and for alternative *a* we obtain a mean p*K*_a_-value in coupling frames of 6.4 ± 0.7 (Fig. 11d). If E285 is protonated, the coupling to D308 in alternative *b* yields a mean p*K*_a_-value of 6.2 ± 0.6 for E285, and the corresponding distribution of all frames is almost centered on the critical region between pH 6 and 7 (Fig. 11b). The effect is not as strong, if E293 is protonated (Fig. 11b). For alternative *a* the p*K*_a_-distribution of D308 reaches slightly into the region between pH 6 and 7, and for alternative *b* the p*K*_a_- distribution of E293 reaches into this region. However, the corresponding p*K*_a_-values, 5.2 ± 0.7 and 5.5 ± 0.7, are clearly lower than those for the coupling between D308 and E285.

These results show that in the absence of Ca^2+^, D308 and E285 can form a protonated dyad with an effective p*K*_a_ that is likely high enough to sense a pH change from 6 to 7. We therefore believe that the second pH sensor that is active in the H294A mutant is the dyad between D308 and E285. In wild-type langerin the pH sensor H294 and this dyad would amplify each other: the K257–D308 hydrogen bond increases the probability that D308 is protonated, and, after Ca^2+^ has escaped, the protonated D308 is stabilized by the D308–E285 dyad. Constant-pH simulations (61–63) or QM/MM simulations (64, 65) could be used to verify whether D308 and E285 indeed form a dyad and constitute the second pH sensor.

Note that the conformational fluctuations in the Ca^2+^- binding pocket give rise to large fluctuations in the instantaneous p*K*_a_-value (Fig. 11) with some distributions covering more than six p*K*_a_ units. Thus, knowing the underlying conformational distribution is essential for a reliable estimate of the overall p*K*_a_-value.

### Comparison to other C-type lectins

To gain insight into whether the proposed mechanism for the pH sensitivity is found in other C-type lectins, we compared the sequences of human langerin to mouse langerin and to human variants of 15 related C-type lectins (SI Fig. 31). All 16 proteins exhibit the typical C-type-lectin fold, as evidenced by crystal structures (SI Fig. 32). The residues D308 and E285, which form the proposed second pH sensor, are highly conserved. However, one should be careful to interpret this as evidence for a conserved second pH-sensor, because these residues are also essential for the coordination of Ca^2+^ and might be conserved for this reason.

The H294–K257 motif, the primary pH sensor in langerin, is not conserved in our sequence alignment. Thus, the proposed mechanism for the pH sensitivity of the Ca^2+^ affinity via H294 protonation does not seem to be the most widespread mechanism to sense a change in the environment in C-type lectins. But the sequence alignment points to possible other mechanisms for sensing a change in the environment.

The selectines P-, E-, and LSECtin share the lysine K257 with langerin in the same position. Additionally, the preceding threonine T256 in langerin is replaced by an arginine in these three proteins, while H294 is replaced by an aspartic acid. Note that in LSECtin Ca^2+^-affinity increases if the pH decreases (27). It is however unclear whether this reversed pH sensitivity is brought about by the change of the H294–K257 motif. Other lysine residues in the short-loop in comparable positions as K257 in langerin can also be found in MCL, lung surfactant protein (SP), CD23a, Endo180, and MMR.

H294 only appears in human and mouse langerin, and is replaced by aspartic acid in most of the other C-type lectins. Instead we find a Ca^2+^-cation in the position where langerin has the H294–K257 hydrogen bond in 6 out of 15 lectins in our analysis (ASGPR, MBP, DCSIGN, DCSIGNR, SP, SR). This Ca^2+^-cation would be partially solvent exposed even when a large entity (such as a pathogen) is bound to the C-type lectin (SI Fig. 32). One therefore might speculate that these lectins do not sense a change in pH but rather a sense in Ca^2+^ concentration.

Several C-type lectins have histidines in other positions close to the Ca^2+^ binding site, which might act as pH sensors via a different mechanism. As already mentioned, ASGPR has a histidine residue that is close to the Ca^2+^ in the primary Ca^2+^ binding site and thereby acts as pH sensor. Furthermore, dectin-2 and MMR have a histidine residue as a neighbor to a Ca^2+^-coordinating residue, and Endo180 and MCL have histidines at the beginning of the long-loop. Whether these histidines act as pH sensors can be tested by mutating the histidine residue and measuring the pH-sensitivity of the Ca^2+^ affinity and of the carbohydrate affinity. Once a residue is a confirmed as a pH sensor, the approach presented in this contribution can be used to propose a molecular mechanism for the pH sensitivity.

## Conclusion

We have described the consequences of a H294 protonation in langerin and its implications for its biological function as an endocytic pattern recognition receptor. When langerin enters the acidic environment of an endosome, it releases its Ca^2+^ co-factor and subsequently its pathogenic cargo, triggered by a moderate change in pH. The Ca^2+^-binding site is blocked from direct solvent access by the pathogen, and additionally the Ca^2+^-coordinating residues have low protonation probabilities in the presence of calcium. Instead, H294 acts as an accessible site, sensing already a change in pH from 7 to 6 (18).

In this contribution, we have uncovered a mechanism in which protonation of H294 perturbs the hydrogen bonded network of the surrounding residues, and alters the conformational ensemble of langerin. A new conformation becomes accessible, in which the protonated K257 side-chain forms a hydrogen bond with the Ca^2+^-coordinating D308, thereby moving a positive charge into the vicinity of the Ca^2+^-binding site. This alone can facilitate the Ca^2+^- release as shown by the reduction in the required force to pull out the ion from its binding site in our steered MD experiments.

The close availability of K257 as a proton source next to the Ca^2+^-binding site possibly results in a proton transfer to the side-chain of D308. At least it has been shown in a theoretical model, that the neutral form of a lysine-aspartate pair can be favored over the salt-bridge, if the dielectric constant of the medium is low as it can be the case in the environment of a protein (43). Thus, protonation of the initial pH sensor H294 likely triggers a cascade of events that ensures the unbinding of Ca^2+^: K257 transfers a proton to D308, protonation of D308 competes drastically with Ca^2+^-binding and, after Ca^2+^ is expelled, the protonation of D308 is stabilized by a dyad with E285. Protonation of D308 additionally accelerates the unfolding of the long-loop, preventing Ca^2+^ from rebinding.

For langerin’s role as endocytic pattern recognition receptor a fast and irreversible Ca^2+^ release is essential. On the cell surface, Ca^2+^ needs to be tightly bound such that the receptor is continuously ready to bind to pathogens. Yet, after endocytosis langerin is probably recycled within minutes (13, 66). This leaves little time for the release of the pathogen, which must be preceded by the unbinding of Ca^2+^. The mechanism that we proposed is an elegant solution to these contradicting requirements: the Ca^2+^-unbinding rate is increased by the K257–D308 hydrogen bond, and after the initial Ca^2+^ release, a transfer of the proton from K257 to D308 triggers a transition to a conformation to which Ca^2+^ cannot rebind.

Note that while our results show that the K257–D308 interaction decreases the stability of Ca^2+^ in the binding pocket and that the protonation of D308 triggers the long-loop unfolding, the transfer of a proton from K257 to D308 is currently an assumption. More work is needed to study the equilibrium between the initial and the end state of the proton transfer. Computationally, this could be tackled by QM/MM calculations (64, 65), free-energy calculations with classical force fields, or by constant pH simulations (61–63).

Another concern is that the point charge Ca^2+^ model might not capture the energetics of Ca^2+^-binding accurately enough, because the point charge model does not enforce coordination and neglects polarization effects. In our study, we tried to minimize the influence of these force field effects by analyzing the differences between two protonation states. However, more elaborate Ca^2+^ models such as reparametrized point-charge models (67, 68), multisite models (69), or polarizable models (70) are available, and should be used for example for the computation of state-specific Ca^2+^ binding free-energies.

Our close atomistic inspection of langerin and its conformational shift upon protonation, gives insight into how pH-sensitivity can be incorporated in biological systems. What seemed like a general conformational shift upon protonation in Fig. 3 could be focused to a specific rearrangement of a side chain (K257) to transport the information from the primary pH sensor (H294) to the allosterically regulated site (Ca^2+^-binding site). Even though the H294–K257 motif is not typical for C-type lectins, many of these proteins exhibit a highly specific pH sensitivity and have potential pH sensors in the vicinity of the primary Ca^2+^-binding site. Our approach can serve as a road map to elucidate the mechanism of pH sensitivity in these systems.

## Experimental procedures

### Molecular dynamics simulations

We used the software package GROMACS (71–77) in setup and production to simulate the considered systems in the *NPT*-ensemble (1 bar, 300 K) with AMBER99SB-ILDN force-field parameters (78) and the TIP3P water model (79). Prior to production, starting structures were put into a sufficiently large simulation box, solvated, neutralized and equilibrated for several hundred picoseconds. For further details refer to the SI.

### Protein expression and purification

All standard chemicals and buffers used within this work were purchased from Sigma Aldrich (St. Louis, MO, USA) or Carl Roth (Karlsruhe, Germany) if not indicated otherwise.

Human langerin CRD WT and all mutants (amino acids 193-328) were cloned from a codon-optimized langerin gene for bacterial expression (GenScript, Piscataway, NJ, USA) into a pET-28a vector (GenScript, Piscataway, NJ, USA) with His-tag, T7 promoter and Kanamycin resistance. Insoluble expression was performed in *E. coli* BL21 (ThermoFisher Scienific, Waltham, MA, USA) in LB medium or in isotope-labeled M9 medium at 37 °C. Protein expression was induced by adding 1 mM IPTG. Bacteria were harvested 3-4 h after induction by centrifugation at 4.000 g for 30 min. Cell pellets were lysed in lysis buffer (50 mM Tris, 150 mM NaCl, 10 mM MgCl2, 0.1 % Tween-20, Ph 8) with 1 mg mL^−1^ lysozyme and 100 µg mL^−1^ DNase I (Applichem, Darmstadt, Germany) for at least 3 h at RT. Inclusion bodies were washed twice with 20-30 mL lysis buffer and twice with water to remove soluble proteins.

Inclusion bodies were denatured in 20 mL of denaturation buffer (6 M guanidinium hydrochloride in 100 mM Tris, pH 8) with 0.01 % β-mercaptoethanol for at least 1 h at 37 °C by shaking or overnight at 4 °C by rotating. After centrifuging (15.000 g, 90 min, 4 °C), the supernatant was slowly diluted 1:10 with langerin refolding buffer (0.4 M L-arginine in 50 mM Tris, 20 mM NaCl, 0.8 mM KCl, pH with 1 mM reduced glutathion (GSH) and 0.1 mM oxidized glutathion (GSSG) while stirring at 4 °C for at least 24 h. The refolded protein solution was dialyzed against 2 L TBS buffer (50 mM Tris, 150 mM NaCl, 5 mM CaCl2) and subsequently centrifuged to remove precipitated protein (15.000 g, 90 min, 4 °C). The supernatant was purified via Ni-NTA agarose affinity chromatography and the elution fractions were pooled and dialyzed against MES (25 mM MES, 40 mM NaCl, pH 6) or HBS (25 mM HEPES, 150 mM NaCl, pH 7) buffer. Precipitated protein was removed by centrifugation (15.000 g, 90 min, 4 °C) and the supernatant was used for experiments. Note that this procedure varies slightly from the one in our previous paper (18).

### ITC measurements

Isothermal titration calorimetry experiments were performed using a MicroCal iTC200 (Malvern Instruments, Malvern, UK) using either chelex-filtered HBS (25 mM HEPES, 150 mM NaCl, pH 7) or low salt MES buffer (25 mM MES, 40 mM NaCl, pH 6). The titrant was dissolved in the same buffer as was used for dialysis of the protein sample. Using the iTC200, the titrant CaCl_2_ (15 mM stock) was added in defined steps of 1-2.5 µL to 80 µL protein solution at 298 K while stirring at 750 rpm. The differential heat of each injection was measured and plotted against the molar ratio. The data was fitted to a one-set of sites binding model assuming a Hill coefficient of 1. Due to the low c-values of the measurements (c < 5), the enthalpy could not be determined reliably. See also SI Fig. 29 and 30.

## Acknowledgment

Funded by the Deutsche Forschungsgemeinschaft (DFG, German Research Foundation) under Germany’s Excellence Strategy – EXC 2008 – 390540038 – UniSysCat. The authors thank the North-German Supercomputing Alliance (HLRN), the Paderborn Center for Parallel Computing PC^2^ and the ZEDAT of the FU Berlin for computing time. Also funded by the Deutsche Forschungsgemeinschaft (DFG, German Research Foundation) through CRC 765, and DFG RA1944/6-1. We thank the Max Planck Society for support.

