## Supplemental Information for "Molecular mechanism of the pH-dependent calcium affinity in langerin"

**Molecular mechanism of the pH-dependent calcium affinity in langerin – Supporting information**

### Molecular dynamics simulations

With small deviations in the individual setup procedure, all considered systems have been prepared according to the same general protocol. Langerins carbohydrate recognition domain (CRD) consisting of the residues G198 to P325 is the basis for all simulations. Initial starting conformations were derived from PDB crystal structures of holo-langerin (3p5h, 3p5g or 3kqg) (1, 2) by removing crystal water, ligands and excessive residues and ions as needed. Further starting structures were manually selected from completed simulations focusing on rarely visited conformations. GROMACS 2016+ (3–9) was used for the subsequent tasks. Desired protonation states of the protein were automatically attained during topology preparation. As force field for the protein and the $Ca^{2+}$-cofactor AMBER99SB-ILDN (10) was chosen. The system was put into a dodecahedral box at a distance of at least 1 nm to the box borders, solvated in explicit TIP3P water (11) and minimized (steepest decent, eps $< 1000\,\mathrm{kJ\,mol^{-1}\,nm^{-1}}$) after the replacement of water by chloride to neutralize contingent charges. Equilibration was done with position restraints on all heavy atoms of the solute in the *NVT*- (300 K, V-rescale thermostat (12), coupling rate at 0.1 ps, coupling groups protein and non-protein, 100 ps length) and *NPT*-ensemble (1 bar, Parrinello-Rahman barostat (13), isotropic, coupling rate at 2 ps, 150 ps length). For both, equilibration and production, the LINCS (14) (order 4, 1 iteration) algorithm was applied to constrain all bonds. The leap-frog integrator (15) was used at a time step of 2 fs. For Lennard-Jones (cut-off) and electrostatic (PME (16), order 4) interactions the cut-off was set to at least 1 nm, while the Verlet cut-off scheme was used to create the neighbor lists. Periodic boundary conditions were imposed in all three dimensions. Protein/calcium coordinates were written to a compressed trajectory file at a time step of 1 ps. The individual length of the production simulations varies between 0.15 and 1 µs as listed in the respective tables below (see Tab. 2 to 12). In cases where the total number of replicas and the number of starting structures do not match, multiple simulations have been started from the same structure with different initial velocities.

#### System nomenclature

To distinguish the langerin systems under consideration we denote the cofactor bound state with one letter: a – apo, h – holo. For the protonation state we use a summation scheme in which each protonation site contributes a specific increment. For example, the H229 protonation adds the value 1, while the H294 protonation adds the value 2. If both protonations are present, this is indicated by the sum (1 + 2, e.g. in h3). Refer to table 1 for an overview. Finally we numerate the different mutations, e.g. m1 for H294A. Later we also differentiate the folding state of the long-loop in italic: *f* – folded, *uf* – unfolded. Similarly we mark the key conformation of the H294 protonated systems where the K257–D308 and the H294–E261 hydrogen bonds are formed (quoted as the "green" PCA-cluster in the main document) with an asterisk ($^*$).

Table 1: Langerin system state notation scheme.

| bound | protonation | mutation | conformation |
|---|---|---|---|
| h: holo | H229: 1 | m1: H294A | *f*: folded |
| a: apo | H294: 2 | m2: E261D | *uf*: unfolded |
| | E285: 4 | m3: K257A | $^*$: K257s–D308s/H294s–E261s |
| | E293: 8 | | |
| | D308: 16 | | |

#### Neutral holo-langerin (h0)

In the neutral protein state, the side-chains of the acidic residues aspartic and glutamic acid are modeled as deprotonated (charge −1) and those of the basic residues arginine and lysine as protonated (charge +1). All other residues including histidine are kept neutral. The presence of the bound calcium ion results in a net-charge of the system of +2, which was neutralized by two chloride atoms in solution. For neutral histidine two different tautomers, the δ- and ϵ-state exist. In high pH solutions of the free acid the ratio of the tautomers is ~1 : 4 in favor of the ϵ-form. In a protein environment, however, the ratio depends strongly on stabilization factors like possible hydrogen bond formation (17–19). For each simulation starting structure the corresponding state was automatically determined during topology preparation, always resulting in the ϵ-form for H294. For H229 the used form is denoted in table 2.

Table 2: Simulation overview for h0.

| #Replica | #Starting structures | #δ | #ϵ | $t/\mu s$ | $t_{tot}/\mu s$ |
|---|---|---|---|---|---|
| 1 | 1 | 0 | 1 | 1 | 1 |
| 1 | 1 | 0 | 1 | 0.58 | 0.58 |
| 5[a] | 5 | 5 | 0 | 0.4 | 2 |
| 5 | 1 | 5 | 0 | 0.25 | 1.25 |
| 119 | 14 | 43 | 76 | 0.22 | 26.18 |
| 2 | 1 | 1 | 1 | 0.15 | 0.3 |
| 133 | 23 | 54 | 79 | | 31[b] |

[a] Simulations reused from our previous paper (20), which uses a slightly different preparation protocol. [b] About 1.2 µs of the simulation data contain no calcium output coordinates. Calcium in the binding pocket is present during the simulation but the trajectories cannot be used for analyses that include the calcium positions.

#### Histidine protonated holo-langerin (h3)

In the histidine protonated protein state, the side-chains of H229 and H294 are both modeled as protonated. All other protonation states are as in h0. The presence of the bound calcium ion and the two side-chain protonations results in a net-charge of the system of +4, which was neutralized by four chloride atoms in solution.

Table 3: Simulation overview for h3.

| #Replica | #Starting structures | $t/\mu s$ | $t_{tot}/\mu s$ |
|---|---|---|---|
| 1 | 1 | 1 | 1 |
| 1 | 1 | 0.63 | 0.63 |
| 7 | 1 | 0.25 | 1.75 |
| 104 | 15 | 0.22 | 22.88 |
| 2 | 1 | 0.2 | 0.4 |
| 2 | 1 | 0.15 | 0.3 |
| 117 | 20 | | 27[b] |

[b] About 6.3 µs of the simulation data contain no calcium output coordinates. Calcium in the binding pocket is present during the simulation but the trajectories cannot be used for analyses that include the calcium positions.

#### Neutral holo-langerin H294A mutant (h0m1)

The H294A mutant was created in silico using the VMD (21) molefacture plugin from the crystal structure. The presence of the bound calcium ion results in a net-charge of the system of +2, which was neutralized by two chloride atoms in solution.

Table 4: Simulation overview for h0m1.

| #Replica | #Starting structures | $t/\mu s$ | $t_{tot}/\mu s$ |
|---|---|---|---|
| 10 | 10 | 0.25 | 2.5[a] |
| 20 | 1 | 0.25 | 5 |
| 30 | 11 | | 7.5 |

[a] Simulations reused from our previous paper (20), which uses a slightly different preparation protocol.

##### Neutral apo-langerin (a0)

The neutral apo-state was prepared in the same way as the neutral holo-state after manual deletion of the $Ca^{2+}$-ion from the structure. Note that in all simulations H229 is modeled in its δ-form, while H294 always remains in its ϵ-form.

Table 5: Simulation overview for a0.

| #Replica | #Starting structures | $t/\mu s$ | $t_{tot}/\mu s$ |
|---|---|---|---|
| 1 | 1 | 1 | 1 |
| 6 | 2 | 0.5 | 3.0 |
| 4 | 1 | 0.25 | 1 |
| 22 | 2 | 0.22 | 4.84 |
| 4 | 1 | 0.1 | 0.4 |
| 37 | 7 | | 10 |

##### Histidine protonated apo-langerin (a3)

The protonated apo-state was prepared in the same way as the protonated holo-state after manual deletion of the $Ca^{2+}$-ion from the structure. The presence of two protonations results in a net-charge of the system of +2, which was neutralized by two chloride atoms in solution.

Table 6: Simulation overview for a3.

| #Replica | #Starting structures | $t/\mu s$ | $t_{tot}/\mu s$ |
|---|---|---|---|
| 1 | 1 | 1 | 1 |
| 1 | 1 | 0.5 | 0.5 |
| 6 | 1 | 0.25 | 1.5 |
| 51 | 8 | 0.22 | 11.22 |
| 1 | 1 | 0.15 | 0.15 |
| 60 | 12 | | 14 |

##### Histidine and glutamic acid 285 protonated apo-langerin (a6, a7)

The apo-state a6 is protonated at H294 and E285. The presence of two protonations results in a net-charge of the system of +2, which was neutralized by two chloride atoms in solution.

Table 7: Simulation overview for a6.

| #Replica | #Starting structures | $t/\mu s$ | $t_{tot}/\mu s$ |
|---|---|---|---|
| 4 | 3 | 0.25 | 1 |

The apo-state a7 is protonated at H229, H294 and E285. The presence of three protonations results in a net-charge of the system of +3, which was neutralized by three chloride atoms in solution.

Table 8: Simulation overview for a7.

| #Replica | #Starting structures | $t/\mu s$ | $t_{tot}/\mu s$ |
|---|---|---|---|
| 22 | 6 | 0.22 | 4.84 |
| 4 | 1 | 0.21 | 0.84 |
| 1 | 1 | 0.18 | 0.18 |
| 3 | 2 | 0.17 | 0.51 |
| 30 | 6 | | 6.37 |

##### Histidine and glutamic acid 293 protonated apo-langerin (a10, a11)

The apo-state a10 is protonated at H294 and E293. The presence of two protonations results in a net-charge of the system of +2, which was neutralized by two chloride atoms in solution.

Table 9: Simulation overview for a10.

| #Replica | #Starting structures | $t/\mu s$ | $t_{tot}/\mu s$ |
|---|---|---|---|
| 4 | 4 | 0.25 | 1 |

The apo-state a11 is protonated at H229, H294 and E293. The presence of three protonations results in a net-charge of the system of +3, which was neutralized by three chloride atoms in solution.

Table 10: Simulation overview for a11.

| #Replica | #Starting structures | $t/\mu s$ | $t_{tot}/\mu s$ |
|---|---|---|---|
| 1 | 1 | 0.50 | 0.50 |
| 28 | 6 | 0.22 | 6.16 |
| 1 | 1 | 0.21 | 0.21 |
| 1 | 1 | 0.17 | 0.17 |
| 3 | 3 | 0.16 | 0.48 |
| 2 | 2 | 0.15 | 0.30 |
| 5 | 2 | 0.14 | 0.70 |
| 1 | 1 | 0.13 | 0.13 |
| 1 | 1 | 0.10 | 0.10 |
| 43 | 6 | | 8.75 |

##### Histidine and aspartic acid 308 protonated apo-langerin (a18, a19)

The apo-state a18 is protonated at H294 and D308. The presence of two protonations results in a net-charge of the system of +2, which was neutralized by two chloride atoms in solution.

Table 11: Simulation overview for a18.

| #Replica | #Starting structures | $t/\mu s$ | $t_{tot}/\mu s$ |
|---|---|---|---|
| 4 | 4 | 0.25 | 1 |

The apo-state a19 is protonated at H229, H294 and D308. The presence of three protonations results in a net-charge of the system of +3, which was neutralized by three chloride atoms in solution.

Table 12: Simulation overview for a19.

| #Replica | #Starting structures | $t/\mu s$ | $t_{tot}/\mu s$ |
|---|---|---|---|
| 2 | 2 | 0.55 | 1.10 |
| 2 | 2 | 0.51 | 1.02 |
| 17 | 6 | 0.22 | 3.74 |
| 2 | 2 | 0.21 | 0.42 |
| 2 | 2 | 0.16 | 0.32 |
| 4 | 2 | 0.15 | 0.60 |
| 1 | 1 | 0.10 | 0.10 |
| 30 | 6 | | 7.30 |

#### DSSP analysis

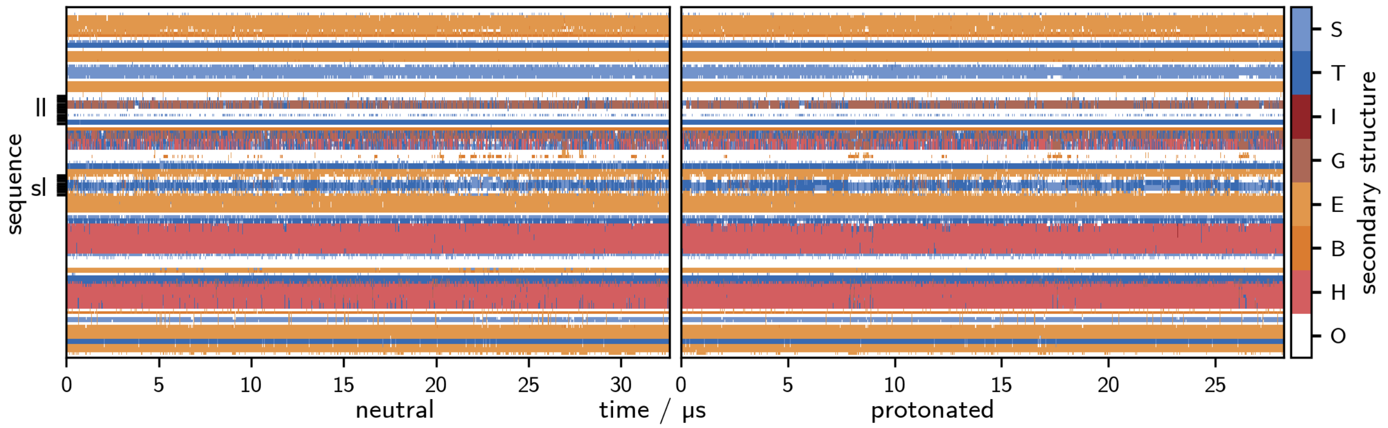


Figure 1: Analysis by the hydrogen bond estimation algorithm DSSP of the secondary structure in the neutral (left) and the protonated holo-state (right). Legend: S – bend, T – hydrogen bonded turn, I – 5-helix, G – 3-helix, E – extended strand, part of β-ladder, B: isolated β-bridge, H: α-helix, O: unassigned.

#### Calculation of p$K_a$-values with PROPKA

For selected conformational snapshots from simulations with a reduced time resolution of 1 ns (table 13), p$K_a$-values of titrable amino acids have been estimated using PROPKA 3.1 (22, 23), allowing to treat groups automatically as coupled where appropriate. D308 was detected in some frames non-covalently coupled to either E285 or E293 or both. In frames with no coupling, there is only one p$K_a$-value per residue. For frames with two coupling residues (dyad), PROPKA provides a pair of p$K_a$-values per residue, a higher value for the situation as proton acceptor and a lower one for the role as supporting nucleophile. We report these pairs in the way that the higher p$K_a$ for D308 and the corresponding lower values for E285 and E293 are treated as alternative. For frames in which E285 and E293 couple to D308 at the same time (triad), PROPKA returns a set of different p$K_a$-values per residue based on interaction permutations. We report the minimum and the maximum of these values as a pair for each residue.

Table 13: Overview p$K_a$-value calculation.

| | | | #coupling conformations | | |
|---|---|---|---|---|---|
| System | #conformations | #folded | D308–E285 | D308–E293 | E285–D308–E293 |
| h0 | 30,192 | 30,192 | 10,684 | 4,003 | 11,175 |
| h3 | 20,795 | 20,795 | 7,341 | 2,094 | 8,842 |
| a0 | 10,363 | 5,310 | 20 | 4 | – |
| a3 | 14,029 | 7,127 | 26 | – | – |
| a7 | 6,368 | 4,435 | 4,744 | – | 1 |
| a11 | 8,702 | 6,315 | 22 | 2,333 | 11 |
| a19 | 7,289 | 2,193 | 4,062 | 72 | 26 |

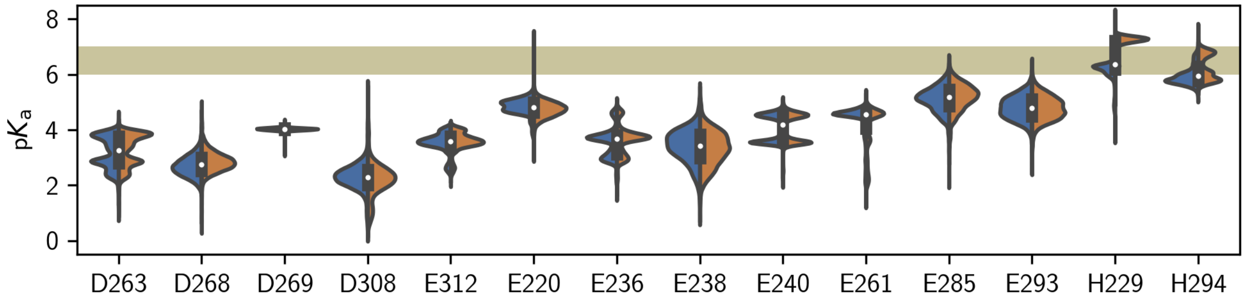


Figure 2: Calculated p$K_a$-values for the neutral (blue, h0) compared to the H294 protonated (orange, h3) holo-system.

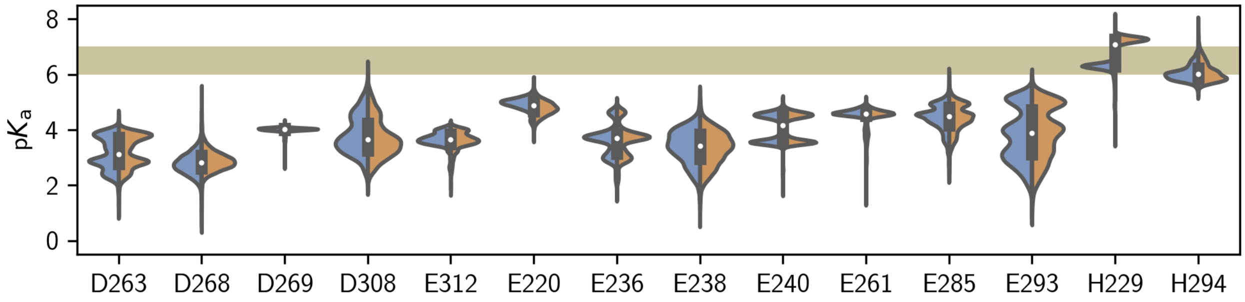


Figure 3: Calculated p$K_a$-values for the neutral (blue, a0) compared to the H294 protonated (orange, a3) apo-system.

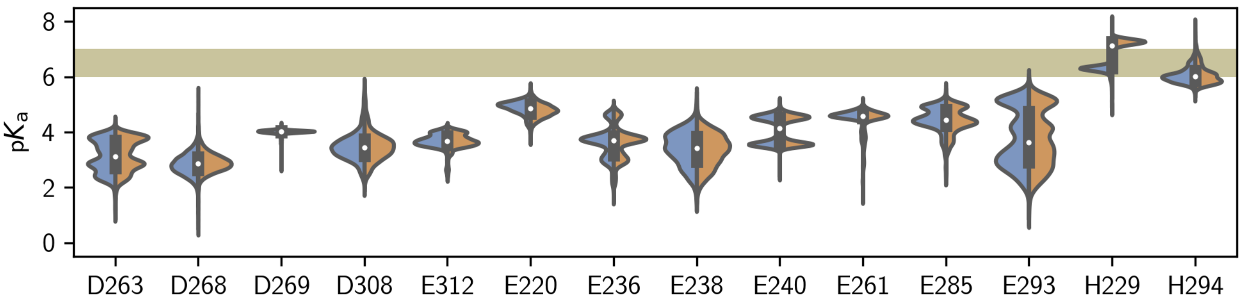


Figure 4: Calculated p$K_a$-values for the neutral (blue, a0) compared to the H294 protonated (orange, a3) apo-system where only folded structures are considered.

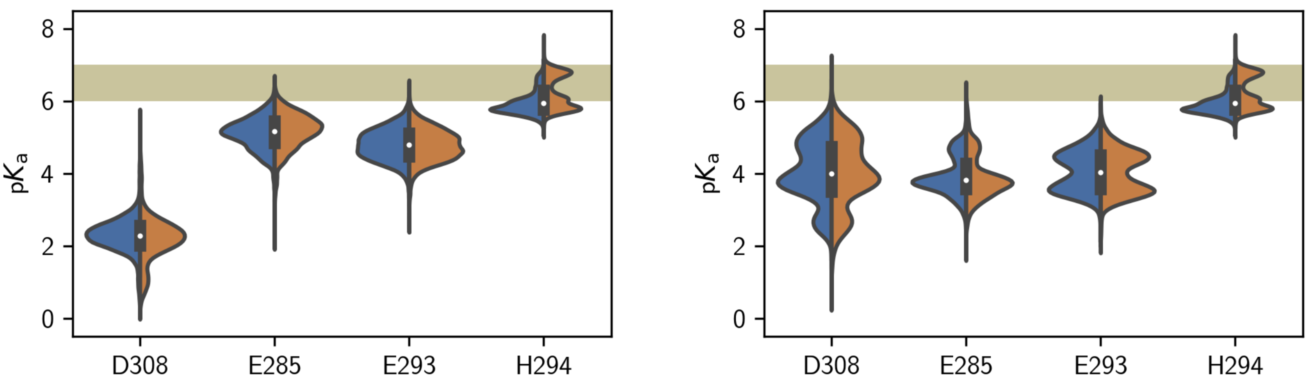


Figure 5: Calculated p$K_a$-values for the neutral (blue, h0) compared to the H294 protonated (orange, h3) holo-system. Alternative p$K_a$-values due to coupling on the right.

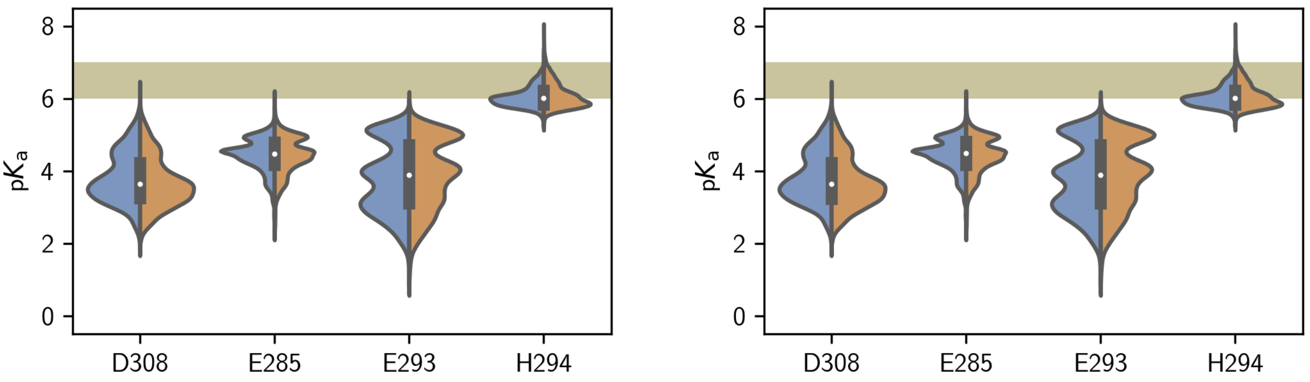


Figure 6: Calculated p$K_a$-values for the neutral (blue, a0) compared to the H294 protonated (orange, a3) apo-system. Alternative p$K_a$-values due to coupling on the right.

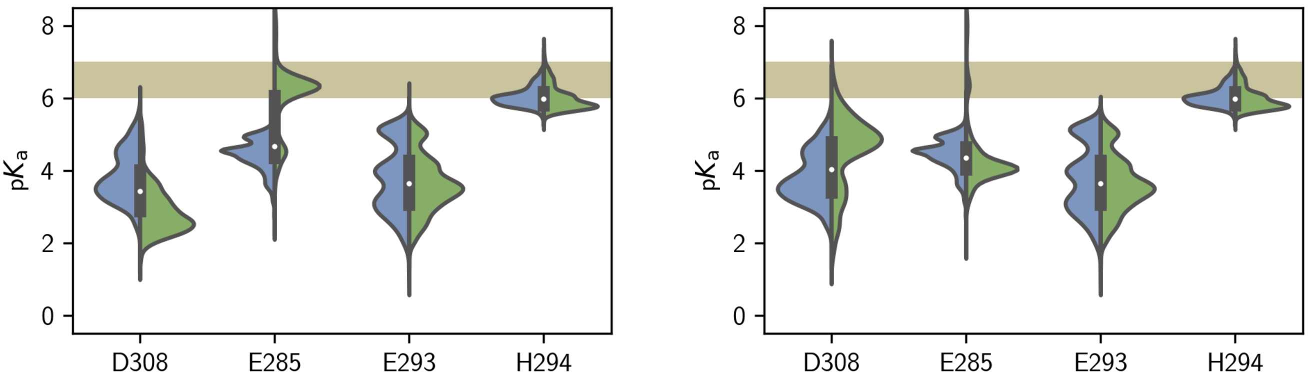


Figure 7: Calculated p$K_a$-values for the neutral (blue, a0) compared to the E285 protonated (green, a7) apo-system. Alternative p$K_a$-values due to coupling on the right.

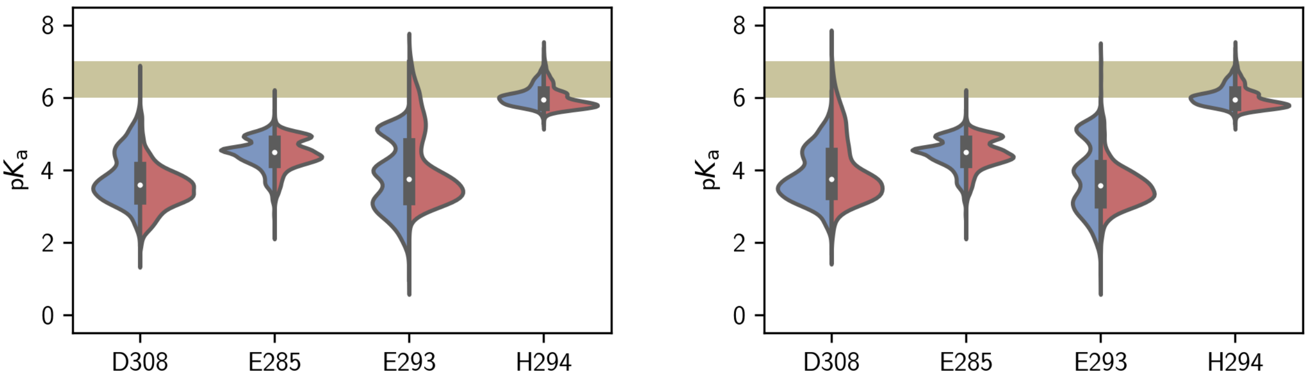


Figure 8: Calculated p$K_a$-values for the neutral (blue, a0) compared to the E293 protonated (red, a11) apo-system. Alternative p$K_a$-values due to coupling on the right.

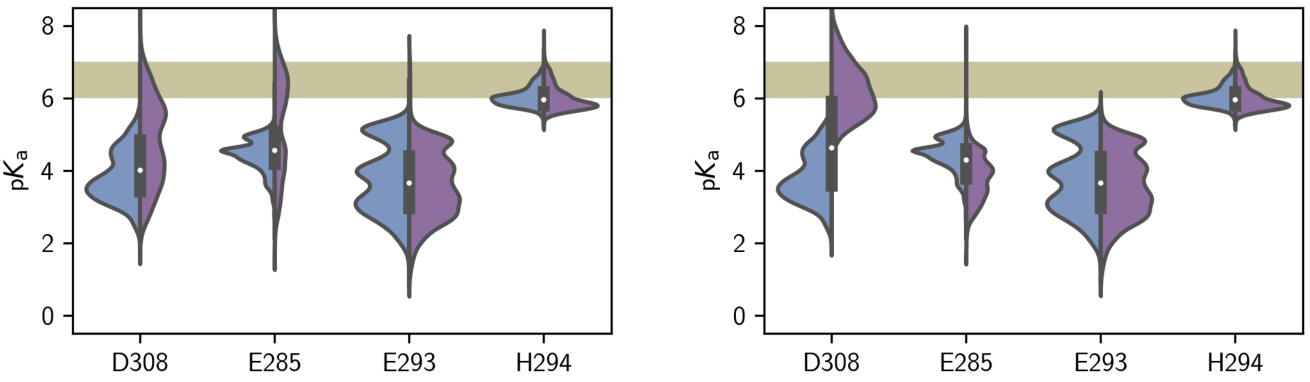


Figure 9: Calculated p$K_a$-values for the neutral (blue, a0) compared to the D308 protonated (purple, a19) apo-system. Alternative p$K_a$-values due to coupling on the right.

Table 14: Mean and standard deviation of calculated p$K_a$-values using PROPKA 3.1 for each ns of simulation time (all frames considered).

| | p$K_a$ $\mu \pm \sigma$ | | | | | | |
|---|---|---|---|---|---|---|---|
| residue | h0 | h3 | a0 | a3 | a7 | a11 | a19 |
| D263 | 3.2 ± 0.6 | 3.3 ± 0.6 | 3.1 ± 0.6 | 3.2 ± 0.6 | 3.1 ± 0.6 | 3.1 ± 0.6 | 3.2 ± 0.6 |
| D268 | 2.7 ± 0.4 | 2.9 ± 0.3 | 2.8 ± 0.4 | 2.9 ± 0.4 | 2.9 ± 0.3 | 2.9 ± 0.3 | 2.9 ± 0.3 |
| D269 | 4.0 ± 0.1 | 4.0 ± 0.0 | 4.0 ± 0.1 | 4.0 ± 0.1 | 4.0 ± 0.0 | 4.0 ± 0.0 | 4.0 ± 0.0 |
| D308 | 2.3 ± 0.5 | 2.2 ± 0.6 | 3.8 ± 0.7 | 3.7 ± 0.7 | 2.9 ± 0.7 | 3.5 ± 0.6 | 4.8 ± 1.2 |
| E312 | 3.5 ± 0.4 | 3.6 ± 0.4 | 3.7 ± 0.3 | 3.6 ± 0.3 | 3.6 ± 0.3 | 3.6 ± 0.3 | 3.6 ± 0.3 |
| E220 | 4.8 ± 0.3 | 4.7 ± 0.3 | 4.9 ± 0.3 | 4.8 ± 0.3 | 4.8 ± 0.3 | 4.8 ± 0.3 | 4.8 ± 0.3 |
| E236 | 3.5 ± 0.5 | 3.6 ± 0.5 | 3.6 ± 0.6 | 3.6 ± 0.6 | 3.6 ± 0.5 | 3.6 ± 0.5 | 3.6 ± 0.5 |
| E238 | 3.4 ± 0.6 | 3.3 ± 0.6 | 3.4 ± 0.6 | 3.3 ± 0.6 | 3.4 ± 0.6 | 3.4 ± 0.6 | 3.4 ± 0.6 |
| E240 | 4.1 ± 0.5 | 4.1 ± 0.5 | 4.1 ± 0.5 | 4.1 ± 0.5 | 4.1 ± 0.5 | 4.1 ± 0.5 | 4.1 ± 0.5 |
| E261 | 4.4 ± 0.6 | 4.0 ± 0.9 | 4.5 ± 0.4 | 4.4 ± 0.5 | 4.5 ± 0.5 | 4.4 ± 0.5 | 4.5 ± 0.3 |
| E285 | 5.1 ± 0.4 | 5.2 ± 0.5 | 4.4 ± 0.5 | 4.4 ± 0.5 | 6.0 ± 0.9 | 4.5 ± 0.4 | 5.2 ± 1.3 |
| E293 | 4.8 ± 0.4 | 4.8 ± 0.4 | 3.8 ± 0.9 | 3.9 ± 0.9 | 3.6 ± 0.8 | 4.1 ± 1.0 | 3.6 ± 0.9 |
| H229 | 6.1 ± 0.4 | 7.1 ± 0.3 | 6.2 ± 0.4 | 7.2 ± 0.3 | 7.2 ± 0.2 | 7.2 ± 0.3 | 7.2 ± 0.3 |
| H294 | 5.9 ± 0.3 | 6.2 ± 0.4 | 6.0 ± 0.3 | 6.1 ± 0.3 | 6.0 ± 0.3 | 6.0 ± 0.3 | 6.0 ± 0.3 |
| | Alternative p$K_a$ $\mu \pm \sigma$ | | | | | | |
| residue | h0 | h3 | a0 | a3 | a7 | a11 | a19 |
| D308 | 4.1 ± 0.9 | 4.0 ± 0.9 | 3.8 ± 0.7 | 3.7 ± 0.7 | 4.5 ± 1.0 | 4.0 ± 0.9 | 6.2 ± 0.8 |
| E285 | 3.9 ± 0.5 | 3.9 ± 0.6 | 4.4 ± 0.5 | 4.4 ± 0.5 | 4.4 ± 0.9 | 4.5 ± 0.4 | 3.8 ± 0.6 |
| E293 | 4.1 ± 0.6 | 4.0 ± 0.6 | 3.8 ± 0.9 | 3.9 ± 0.9 | 3.6 ± 0.8 | 3.6 ± 0.6 | 3.6 ± 0.9 |

Table 15: Mean and standard deviation of calculated burying-values using PROPKA 3.1 for each ns of simulation time.

| | %buried $\mu \pm \sigma$ | | | | | | |
|---|---|---|---|---|---|---|---|
| residue | h0 | h3 | a0 | a3 | a7 | a11 | a19 |
| D263 | 1 ± 2 | 0 ± 1 | 1 ± 2 | 1 ± 3 | 1 ± 2 | 1 ± 2 | 1 ± 2 |
| D268 | 37 ± 13 | 45 ± 11 | 42 ± 10 | 48 ± 8 | 49 ± 6 | 49 ± 6 | 48 ± 8 |
| D269 | 0 ± 0 | 0 ± 0 | 0 ± 0 | 0 ± 0 | 0 ± 0 | 0 ± 0 | 0 ± 0 |
| D308 | 38 ± 4 | 38 ± 4 | 35 ± 6 | 34 ± 5 | 34 ± 5 | 33 ± 5 | 41 ± 7 |
| E312 | 0 ± 0 | 0 ± 0 | 0 ± 0 | 0 ± 0 | 0 ± 0 | 0 ± 0 | 0 ± 0 |
| E220 | 16 ± 7 | 41 ± 10 | 18 ± 7 | 42 ± 10 | 44 ± 7 | 43 ± 9 | 43 ± 9 |
| E236 | 0 ± 0 | 0 ± 0 | 0 ± 0 | 0 ± 0 | 0 ± 0 | 0 ± 0 | 0 ± 0 |
| E238 | 44 ± 10 | 46 ± 8 | 47 ± 8 | 50 ± 8 | 49 ± 7 | 53 ± 10 | 48 ± 8 |
| E240 | 0 ± 0 | 0 ± 0 | 0 ± 0 | 0 ± 0 | 0 ± 0 | 0 ± 0 | 0 ± 0 |
| E261 | 0 ± 0 | 0 ± 0 | 0 ± 0 | 0 ± 0 | 0 ± 0 | 0 ± 0 | 0 ± 0 |
| E285 | 10 ± 7 | 13 ± 8 | 4 ± 5 | 4 ± 6 | 11 ± 11 | 3 ± 5 | 22 ± 12 |
| E293 | 15 ± 4 | 14 ± 4 | 1 ± 3 | 1 ± 3 | 1 ± 3 | 1 ± 3 | 1 ± 3 |
| H229 | 30 ± 13 | 23 ± 5 | 24 ± 12 | 24 ± 5 | 24 ± 4 | 24 ± 4 | 23 ± 4 |
| H294 | 8 ± 9 | 12 ± 9 | 4 ± 6 | 13 ± 9 | 14 ± 9 | 16 ± 8 | 16 ± 8 |

#### Hydrogen bond analysis

We investigated the occurrence of hydrogen bonds in the langerin holo-systems (h0, h3) as we suspected that a change in the protonation state may (transforming a potential hydrogen bond acceptor into a donor) also perturb the observed hydrogen bonded interactions. To get a neutral impression of the present interactions we extracted time-resolved information about all possible bonds, disregarding only those bonds involving the termini (residues 198 to 214 and 322 to 325) and applying an occupancy filter (10 % < occupancy < 90 % in at least one of the systems). The hydrogen bond existence information was collected using the GROMACS tool gmx hbond using the default criteria (hydrogen–donor–acceptor angle $\leq 30°$ and donor-acceptor distance $\leq 3.5$ Å). Figure 10 visualises the resulting subset of interactions sorted by occupancy in h0. Bonds that undergo a large change in occupancy upon H294/H229 protonation (gray highlighting for changes >20 %) can be considered important under the reservation of a potential sampling bias. When a residue of the short- or long-loop is involved, this is indicated by colored circles above the bars. Absolute occupancy is of secondary importance since neither a particularly high nor low value allows a conclusion regarding a contingent function of the structural element.

As a complementary selection criteria we looked for pairwise correlations in the existence of hydrogen bonds, since it can be a hint towards allosteric communication over a hydrogen bonded network. We calculated the Pearson-correlation coefficient for all found (and filtered) hydrogen bonds and refined the result by only looking at those bonds above a correlation cutoff (bond is involved in a correlation >0.6 in at least one of the systems). In figure 10 the resulting correlation matrix is illustrated as a heatmap. The upper left triangle shows the correlation in the individual system while the lower right triangle shows the absolute difference in the correlation between the two systems. The diagonal elements represent again the occupancy of the bonds. Highly correlated bonds are of special interest, in particular if the degree of correlation changes upon protonation.

Combining the decision criteria, occupancy and correlation, and focusing on those bonds formed by residues of the short- and long-loop and respectively the interconnecting segment, we settled on the following set of important hydrogen bonds as structurally interesting main factors effected by histidine H294 protonation: 253m-296m, 256s-294m, 257s-261s, 257s-293m, 257s-294s, 257s-308s, 258m-256m, 262m-259m, 262m-261s, 265s-256m, 266m-270m, 272m-264m, 274s-263s, 276m-273m, 287m-308s, 288s-260m, 288s-261s, 288s-291s, 292m-289m, 294m-292m, 294m-309m, 294s-261s, 294s-291m, 297s-307s, 308m-285s.

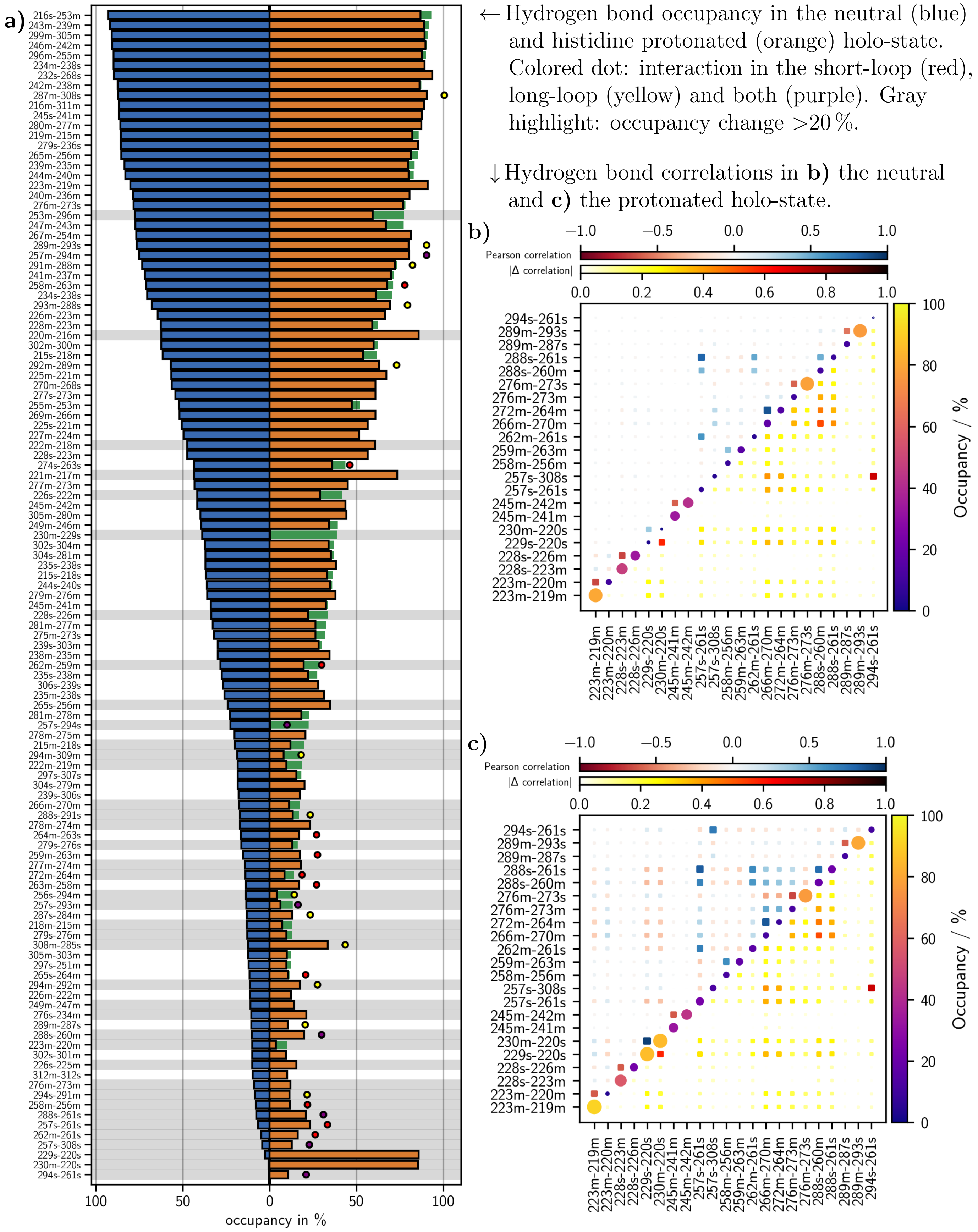


Figure 10: Hydrogen bond analysis.

#### Principle component analysis

For the principle component analysis (PCA) we used the GROMACS tool gmx covar in a joint fashion on a concatenated trajectory of h0 and h3, including only non-hydrogen protein atoms of the residues 201 to 322 (989 atoms), with an increased time step of 1 ns (57481 frames). All structures have been fitted using the backbone atoms of residues 201 to 256 and 295 to 322 (252 atoms) as a reference. Replicawise projections of the individual systems (31091247 frames for h0, 26389202 frames for h3) onto the resulting eigenvectors and structure interpolations have been obtained with the GROMACS tool gmx anaeig.

#### Time-lagged independent component analysis

We used the PyEmma (24) package version 2.5.7 for the time-lagged independent component analysis (TICA). The analysis was done for the neutral (h0) and the protonated (h3) holo-system separately on the same set of input features, which are backbone dihedrals of the residues 253 to 312, $\chi_1$ and $\chi_2$ side-chain dihedrals of the residues 285, 293, 294 and 308 and existence-functions for the set of hydrogen bonds we settled on previously (see section *Hydrogen bond analysis*): 253m-296m, 256s-294m, 257s-261s, 257s-293m, 257s-294s, 257s-308s, 258m-256m, 262m-259m, 262m-261s, 265s-256m, 266m-270m, 272m-264m, 274s-263s, 276m-273m, 287m-308s, 288s-260m, 288s-261s, 288s-291s, 292m-289m, 294m-292m, 294m-309m, 294s-261s, 294s-291m, 297s-307s, 308m-285s. Dihedral angles were decomposed into sine and cosine components. Hydrogen bond existences were converted to trajectories of 1 (present) and -1 (not present). We included 128 replica for the unprotonated and 116 replica for the protonated system with a time-step of 5 ps. Lag times between 1 and 30 ns were tested, giving the most satisfying result with respect to eigenvalue decay and state separation in the projections at 20 ns. Projections in 6D are shown in figure 11.

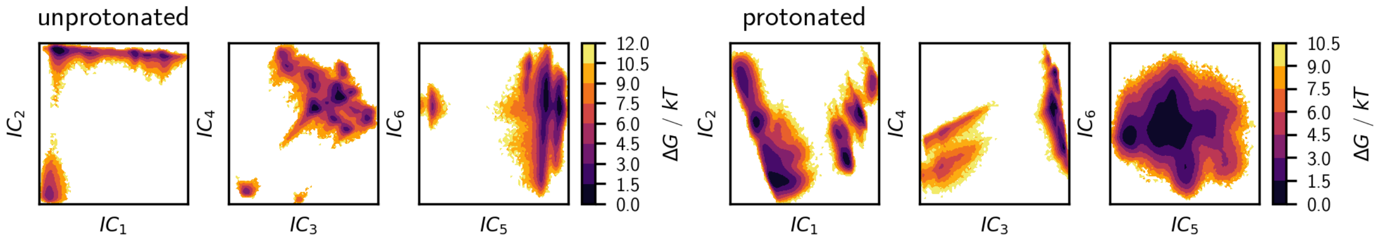


Figure 11: 6D projections of the unprotonated and protonated system onto the first 6 independent components of the TICA.

Figure 12 shows correlations of the input coordinates with components obtained from the TICA to get an impression on what degrees of freedom contribute most to the transformed coordinates.

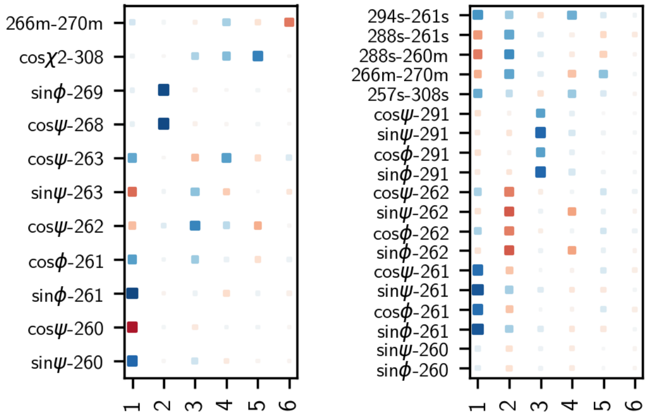


Figure 12: Correlation of input features to the transformed coordinates of the TICA (cutoff >0.5).

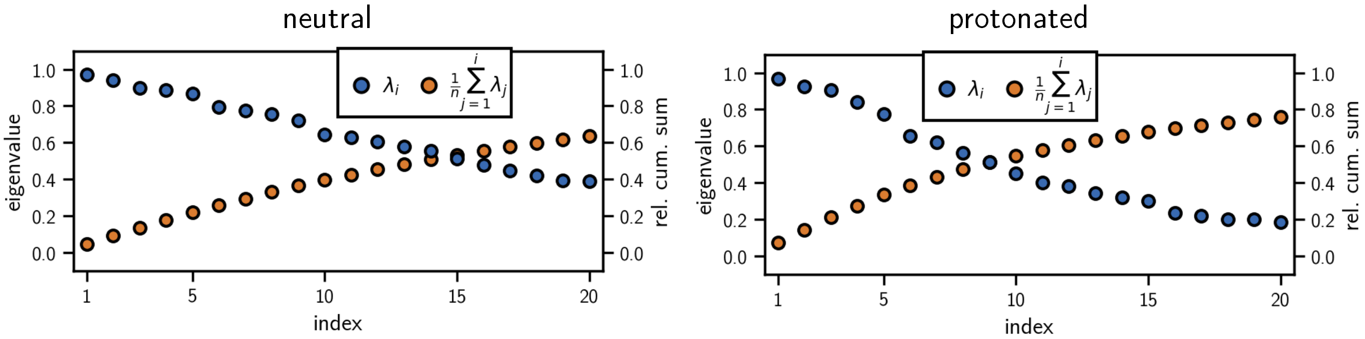


Figure 13: Eigenvalue spectra of the TICA for the neutral and the protonated holo-system.

#### Clustering and core-set Markov-state model construction

For the clustering of data points in the PCA- and TICA-space we used the density-based common-nearest neighbors (CNN) cluster-algorithm in an implementation that is currently under development. The source is available on GitHub: `git@github.com:janjoswig/CNN.git`. The package includes also functionalities for the estimation of core-set Markov-state models on top of a discretization obtained from such a clustering.

In figure 14 we show the final cluster results for the TICA-projections of the neutral and the protonated system. In the unprotonated case we isolated 25 clusters in a 5-step hierarchical procedure. For the protonated system we found 22 clusters in 6 steps. Clustering was done on a reduced data set with a time step of 1 ns. Cluster assignments were subsequently predicted for a data set with a time step of 100 ps.

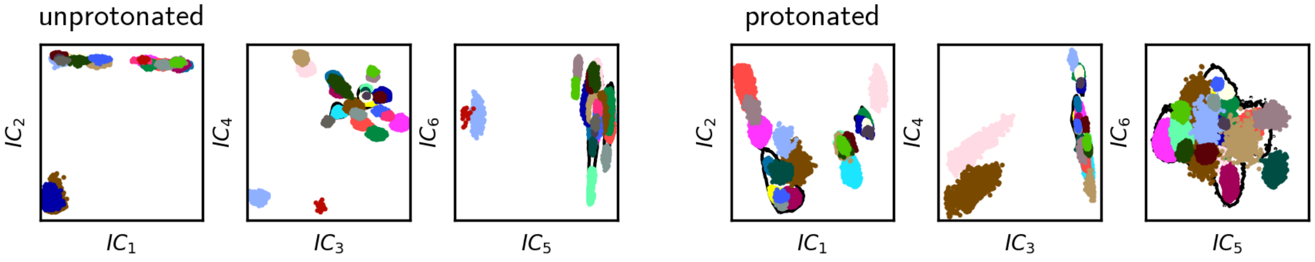


Figure 14: 6D projections of the unprotonated and protonated system onto the first 6 independent components of the TICA clustered into 25 and 22 core-sets, respectively.

We estimated MSMs for lag-times between 1 and 16 ns. Figure 15 shows the implied time-scale teste for these models.

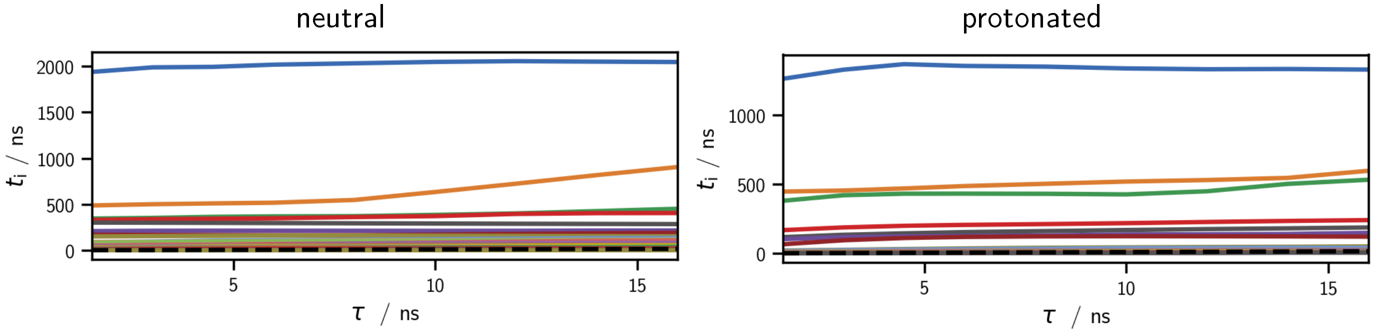


Figure 15: Implied time scales for MSMs of the neutral and the protonated system.

The core-sets were lumped by Perron-cluster cluster analysis (PCCA) on the basis of a MSM with 7 ns lag-time, giving the meta-stable sets of clusters connected by the first 5 transition processes shown in figure 16. Also shown are representative structures for each set of long-lived conformations in figure 17.

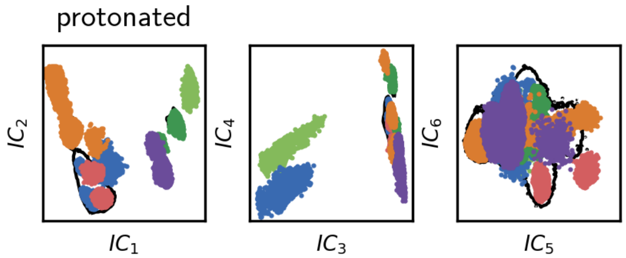


Figure 16: Core-sets from figure 14 grouped to meta-stable sets by PCCA incorporating the 5 slowest processes.

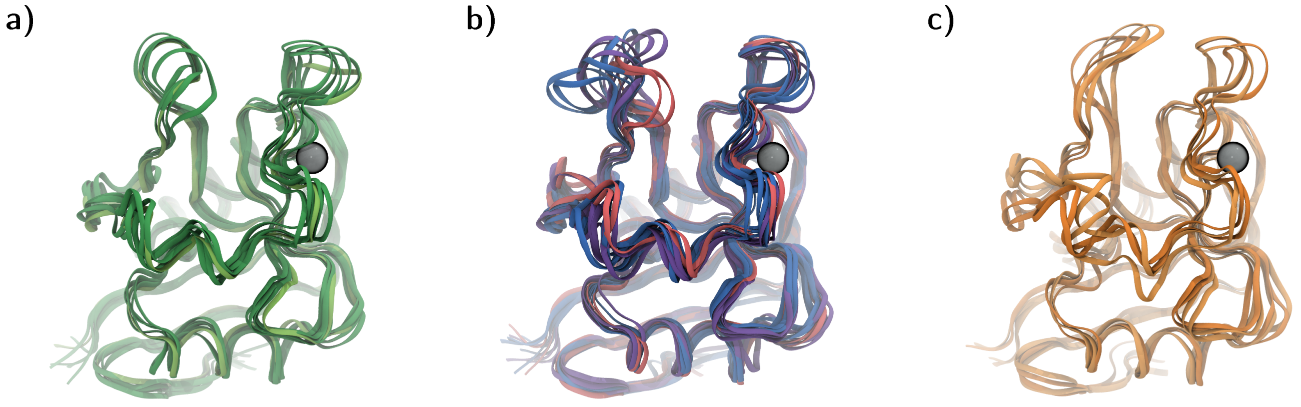


Figure 17: Example structures for meta-stable conformations from figure 16 in the H294-protonated holo-system. **a)** K257s–D308s bonded conformations referred to as the "green" cluster. **b)** Open loop forms with distinct short-loop conformations. **c)** Closed loop-forms referred to as the "orange" cluster. Within this set, $\alpha_3$-helix conformations exchange on a time scale of 130 ns (lighter and deeper orange; not shown in the 2D projection).

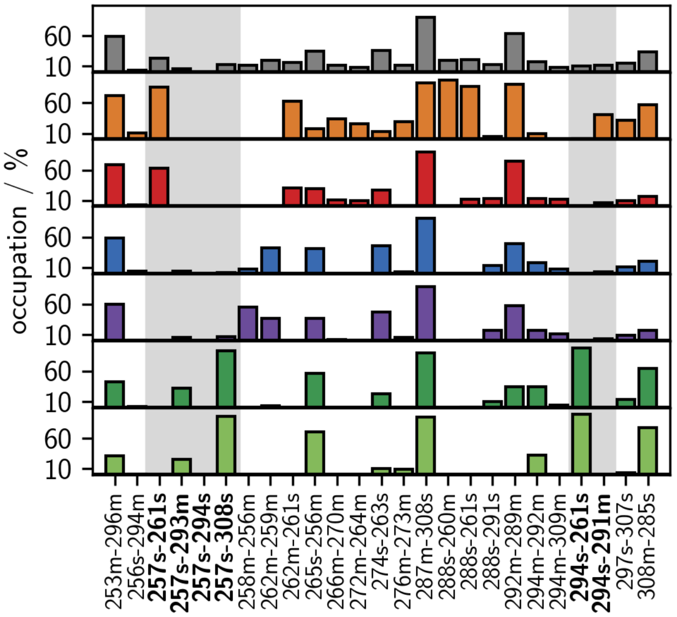


Figure 18: Hydrogen bond occupancy in meta-stable sets of the H294-protonated holo-system from figure 16 (ensemble in gray).

#### K257–D308 distance

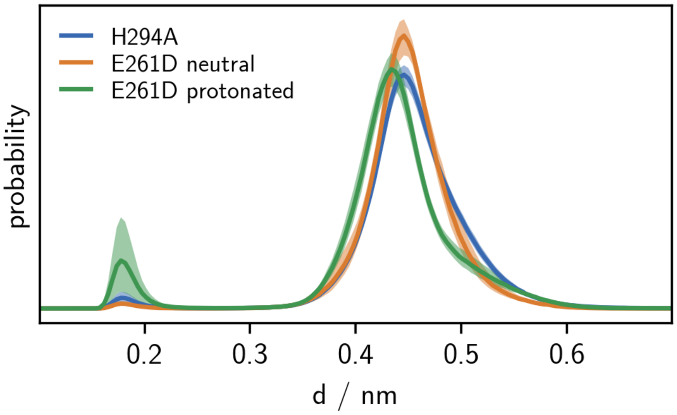


Figure 19: K257–D308 distance distribution for the H294A mutant and E261D with protonated and neutral H294. 95 % confidence interval on the mean over replica-wise probability densities obtained by bootstrapping (1000 samples).

#### Coulomb interactions

Fig. 20 shows the Coulomb force between two positive elementary point-charges as a function of the distance for several typical relative dielectric constants. In our MD simulations with explicit water, the pairwise Coulomb interactions are not truncated and are calculated at a relative dielectric constant of one, i.e following the blue curve in Fig. 20. But the atoms in the space between $Ca^{2+}$ and H294 can rearrange according to the surrounding electrostatic field thereby causing an overall shielding effect, such that the effective force between two residues on the protein surface, e.g. H294 and K257, will be similar to the green curve. Thus, at distances of $d > 1.0$ nm, the effective Coulomb forces between residues on the protein surface are close to zero.

Fig. 21 shows histograms of the pairwise Coulomb-energies in vacuum between H294, K257 and $Ca^{2+}$ of the neutral and H294 protonated holo-state. As expected, the Coulomb energy between $Ca^{2+}$ and the overall neutral H294 is approximately zero, and increases to about 30 kJ mol$^{-1}$ when H294 is protonated. Note that in the presence of water, the corresponding repulsive force is dampened by a shift of the charge distribution in the space between $Ca^{2+}$ and $H294^{+}$. The most striking result of this analysis is the change of interaction energies between $Ca^{2+}$ and K257 upon protonation of H294. In the protonated state the interaction energies reach values beyond 100 kJ mol$^{-1}$. We confirmed that these high energy values correspond to conformations in which the K257s–D308s hydrogen bond is formed, i.e. to those conformations that, according to our model, are responsible for the regulation of the $Ca^{2+}$-affinity. When the K257s–D308s hydrogen bond is formed, the repulsion between the positively charged K257 and $Ca^{2+}$ cannot be mitigated by the rearrangement of atoms in the intervening space. Thus, in these conformations the Coulomb repulsion between K257 and $Ca^{2+}$ is strong enough to explain the observed decrease in $Ca^{2+}$ affinity. Finally, the Coulomb energy between K257 and H294 shifts from negative energy values to positive values upon protonation of H294, in accordance with the observed opening of the hydrogen bond (dotted lines in Fig. 21).

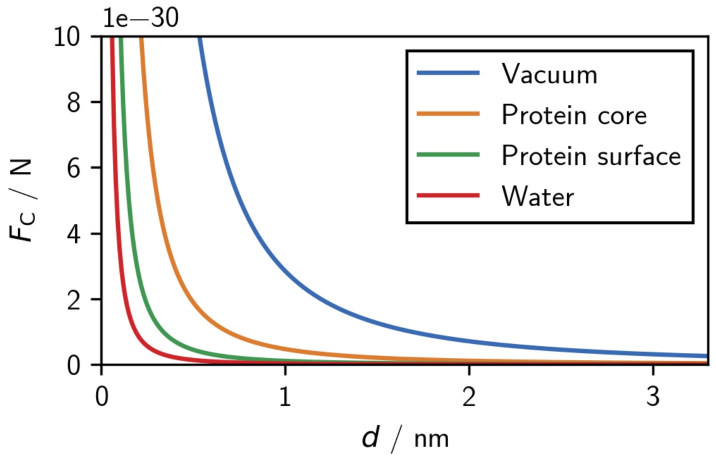


Figure 20: Expected distance dependent Coulomb-force between two positive elementary charges for assumed relative dielectric constants in vacuum ($\epsilon_r = 1$), within a protein ($\epsilon_r = 6$ (25)), on a protein surface ($\epsilon_r = 25$ (25)) and in water ($\epsilon_r = 80$). For media with higher dielectric constants, the interaction is very small beyond distances of 1 nm.

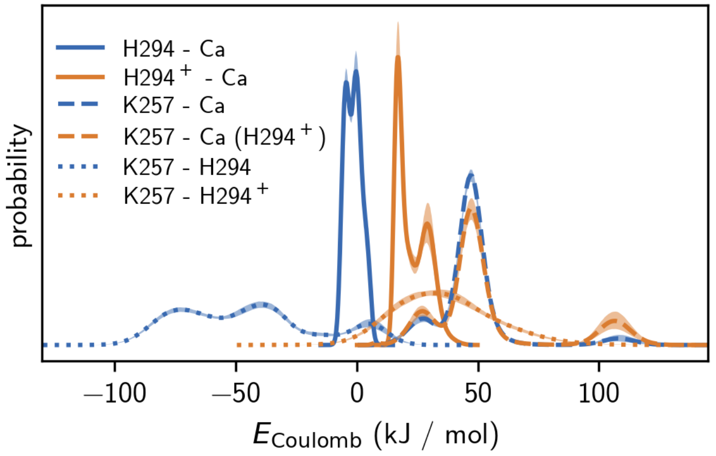


Figure 21: Coulomb contributions during MD runs between H294, K257 and calcium. The interaction energies were obtained from a GROMACS rerun with energy groups on the full trajectory data of neutral (blue) and H294 protonated (orange) holo langerin. 95 % confidence interval on the mean over replica-wise probability densities obtained by bootstrapping (1000 samples).

#### Steered molecular dynamics simulations

All systems simulated in a steered MD setup were prepared following the same general procedure that is very similar to the one used for the conventional MD simulations, but more concise. For a selected starting structure of holo-langerin (as denoted in table 16) without crystal waters, ligands and excessive residues and ions, a topology in the desired protonation state was automatically generated using GROMACS 2019 (3–9) and the AMBER99SB-ILDN force field (10). The molecule was put into a cubic box at a distance of at least 1.5 nm to the box borders and minimized in vacuum (steepest decent, eps $< 100\,\mathrm{kJ\,mol^{-1}\,nm^{-1}}$) before it was solvated in explicit TIP3P water (11) and the replacement of water by chloride to neutralize contingent charges. It was minimized again in solution twice (1. steepest decent, eps $< 100\,\mathrm{kJ\,mol^{-1}\,nm^{-1}}$, 2. conjugate gradient, eps $< 1\,\mathrm{kJ\,mol^{-1}\,nm^{-1}}$) and then equilibrated without any position restraints in the *NVT*- (300 K, V-rescale thermostat (12), coupling rate at 0.1 ps, coupling groups protein and non-protein, 200 ps length) and *NPT*-ensemble (1. 1 bar, Berendsen barostat (26), isotropic, coupling rate at 1 ps, 200 ps length, 2. 1 bar, Parrinello-Rahman barostat (13), isotropic, coupling rate at 2 ps, 400 ps length). For both, equilibration and production, the LINCS (14) (order 6, 2 iterations in equilibration and order 4, 1 iteration in production) algorithm was applied to constrain bonds to hydrogen. The leap-frog integrator (15) was used at a time step of 1 fs during equilibration and 2 fs in production. For Lennard-Jones (cut-off) and electrostatic (PME (16), order 6) interactions the cut-off was set to 1 nm, while the Verlet cut-off scheme was used to create the neighbour lists. Periodic boundary conditions were imposed in all three dimensions. The vacuum minimisation differs substantially from these settings as the group cut-off scheme was utilised, for Lennard-Jones (cut-off) and electrostatic (cut-off) interactions the cut-off was set to infinity (0), and no periodic boundary conditions were applied. In production a force was exerted in form of a harmonic (umbrella) potential along a single coordinate, defined as the distance between the $Ca^{2+}$-ion in the binding pocket and the center of mass of the "upper" protein region represented by the $C_\alpha$-atoms of residues 257, 264, 281, 282, 293 and 294 (see figure 22). The harmonic spring constant was set to $k = 500\,\mathrm{kJ\,mol^{-1}\,nm^{-2}}$ and the pulling-force acting on the pull-coordinate was constantly increased at a velocity of $1 \times 10^{-4}\,\mathrm{nm\,ps^{-1}}$. Protein/calcium coordinates were written to a compressed trajectory file at a time step of 1 ps as well as the instantaneous pull-force. The individual simulations were prolonged to a maximal length of 20 ns or until the pulling-distance exceeded the length of the boxvector. For each system (starting structure) the pull experiment was repeated 40 times. As rupture force, the force needed to remove the $Ca^{2+}$-ion from the binding pocket, we took the maximum pull-force from the force trajectories smoothed by the running mean within a window of 10 ps.

Table 16: Overview steered MD.

| System | Starting structure |
|---|---|
| h0 | crystal structure PDB-ID 3p5g (1) |
| h0*uf* | unfolded conformation selected from a0, calcium inserted manually |
| h0m1 | H294A mutant created with the VMD (21) molefacture plugin, crystal structure analogue |
| h0m3 | K257A mutant created with the VMD (21) molefacture plugin, crystal structure analogue |
| h0m1m3 | K257A/H294A double mutant created with the VMD (21) molefacture plugin, crystal structure analogue |
| h3 | crystal structure analogue |
| h3* | H294–E261/K257–D308 hydrogen bonded conformation selected from h3 ("green" PCA cluster) |
| h3m2* | E261D mutant created with the VMD (21) molefacture plugin, H294–D261/K257–D308 hydrogen bonded conformation |
| h3m3 | K257A mutant created with the VMD (21) molefacture plugin, crystal structure analogue |
| h7 | crystal structure analogue |
| h11 | crystal structure analogue |
| h19 | crystal structure analogue |

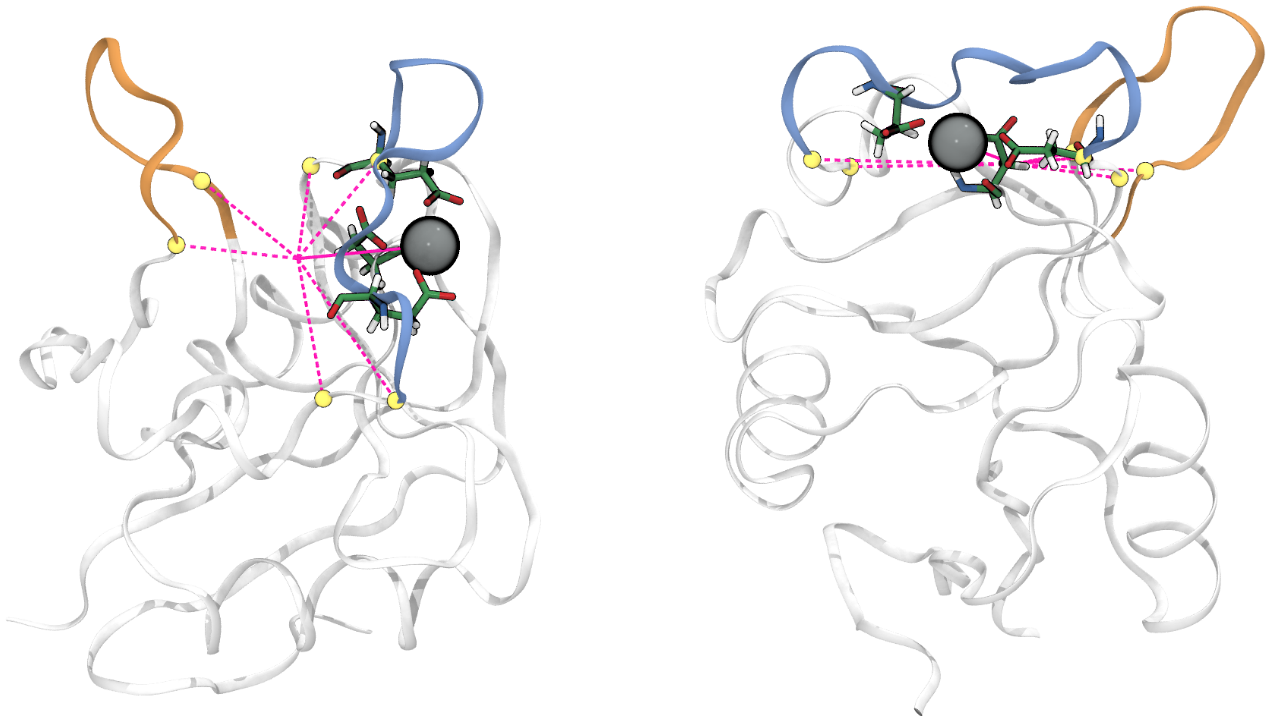

Figure 22: The pull coordinate (solid magenta line) is defined as the distance between the center of mass of the $C_\alpha$-atoms of residues 257, 264, 281, 282, 293 and 294 (yellow spheres, dashed magenta lines), representing the rigid "upper" protein, and the $Ca^{2+}$-ion (gray Van der Waals sphere).

#### Rate determination of apo-langerins long-loop unfolding

Unfolding of apo-langerins long-loop was determined by following five different criteria applied to trajectories with a time step of 1 ns: a) We determined – by visual inspection in VMD (21) – the last frame in which the long-loop appears to be clearly folded. b) If applicable, we also determined the first frame from which on the loop is clearly unfolded irreversibly on the time scale of the simulation. The mean between "last folded" and "first unfolded" point was taken as the unfolding point. c) We tracked the long-loop (residues P283 to E293) $C_\alpha$ root-mean-square deviation (RMSD) with the crystal structure as reference (PDB-ID 3p5g (2)) and detected the first frame in which a value higher than 0.2 nm was reached. d) We tracked the existence of the N287m–D308s hydrogen bond and determined the first frame from which on the bond is not constantly occupied anymore. e) We did the same for N288m–D308s. The hydrogen bond existence functions were obtained as described in section *Hydrogen bond analysis* on page 13. RMSDs were calculated using the GROMACS tool rms. The RMSD time series, smoothed by a running average window of 5 ns, were evaluated by a change-point detection algorithm (mean shift clustering using a bandwidth of 0.035).

Table 17: Trajectories showing an event of unfolding or a clearly folded structure throughout the simulation time of 220 ns by a) visual inspection (last folded frame), b) visual inspection (mean of last folded and first unfolded frame), c) RMSD cut-off (>0.2 nm), d) breaking of N287m–D308s H-bond, e) breaking of N288m–D308s H-bond.

| **a0** | a) | b) | c) | d) | e) |
|---|---|---|---|---|---|
| unfold | 17 | 13 | 15 | 12 | 13 |
| stay folded | 18 | 18 | 20 | 22 | 21 |
| sum | 35 | 31 | 35 | 34 | 34 |
| **a3** | | | | | |
| unfold | 29 | 17 | 29 | 23 | 25 |
| stay folded | 25 | 25 | 25 | 30 | 28 |
| sum | 54 | 42 | 54 | 53 | 53 |
| **a6+a7** | | | | | |
| unfold | 16 | 5 | 6 | 0 | 4 |
| stay folded | 18 | 18 | 24 | 26 | 22 |
| sum | 34 | 23 | 30 | 26 | 26 |

| **a10+a11** | a) | b) | c) | d) | e) |
|---|---|---|---|---|---|
| unfold | 19 | 12 | 13 | 7 | 7 |
| stay folded | 24 | 24 | 26 | 27 | 27 |
| sum | 43 | 36 | 39 | 34 | 34 |
| **a18+a19** | | | | | |
| unfold | 24 | 20 | 19 | 17 | 20 |
| stay folded | 8 | 8 | 9 | 8 | 5 |
| sum | 32 | 28 | 28 | 25 | 25 |

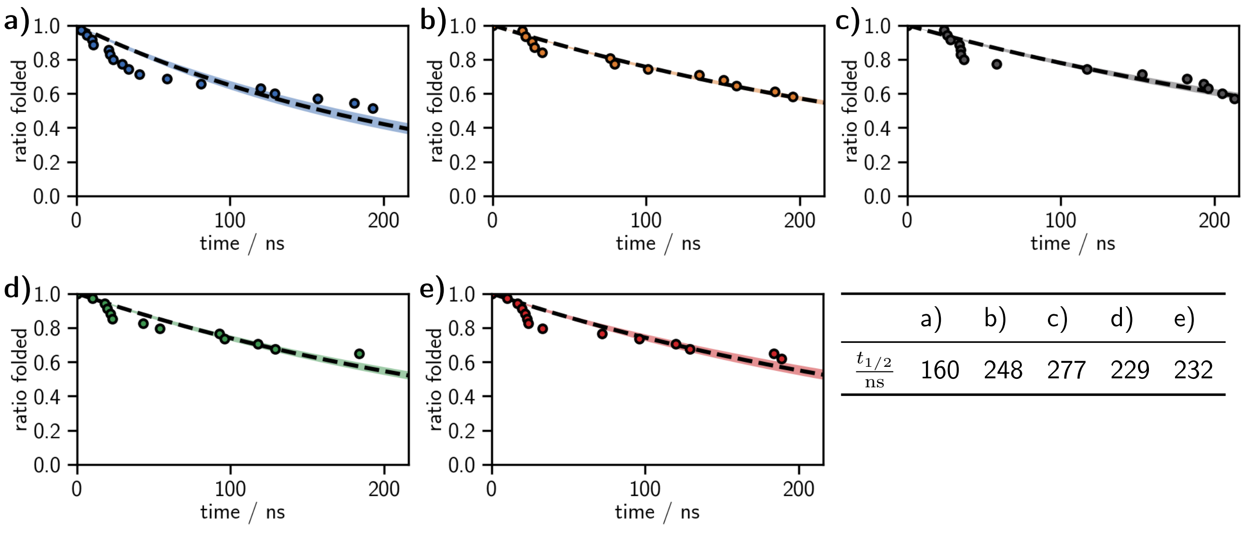


| | a) | b) | c) | d) | e) |
|---|---|---|---|---|---|
| $\frac{t_{1/2}}{\text{ns}}$ | 160 | 248 | 277 | 229 | 232 |

Figure 23: Unfolding events in neutral apo-langerin (a0) and exponential fits measured by **a)** visual inspection (last folded frame), **b)** visual inspection (mean of last folded and first unfolded frame), **c)** RMSD cut-off (>0.2 nm, **d)** breaking of N287m–D308s H-bond, **e)** breaking of N288m–D308s H-bond.

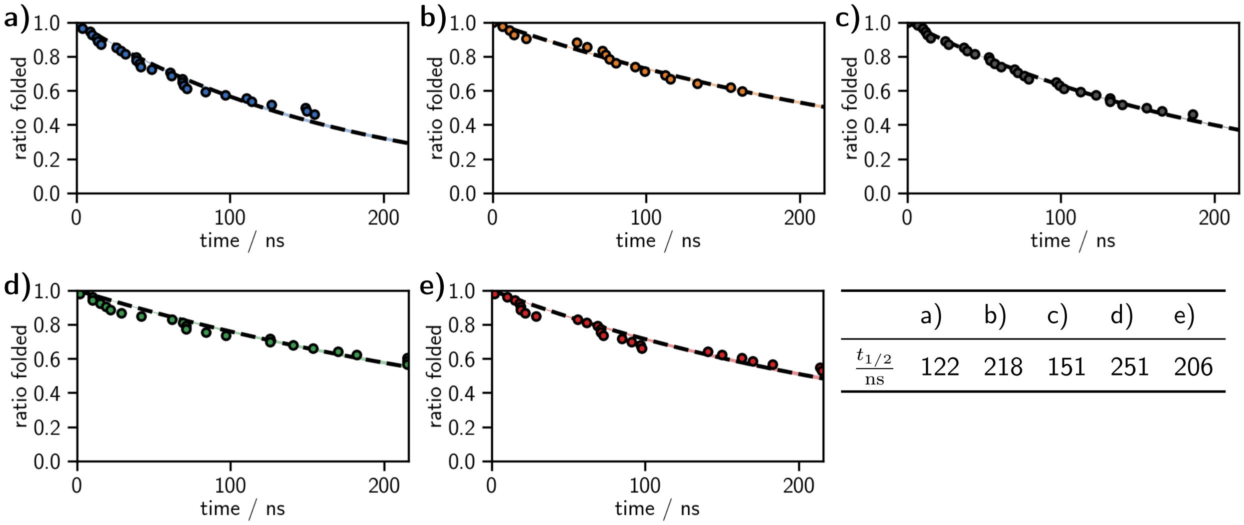

| | a) | b) | c) | d) | e) |
|---|---|---|---|---|---|
| $\frac{t_{1/2}}{\mathrm{ns}}$ | 122 | 218 | 151 | 251 | 206 |

Figure 24: Unfolding events in H294-protonated apo-langerin (a3) and exponential fits measured by **a)** visual inspection (last folded frame), **b)** visual inspection (mean of last folded and first unfolded frame), **c)** RMSD cut-off (>0.2 nm, **d)** breaking of N287m–D308s H-bond, **e)** breaking of N288m–D308s H-bond.

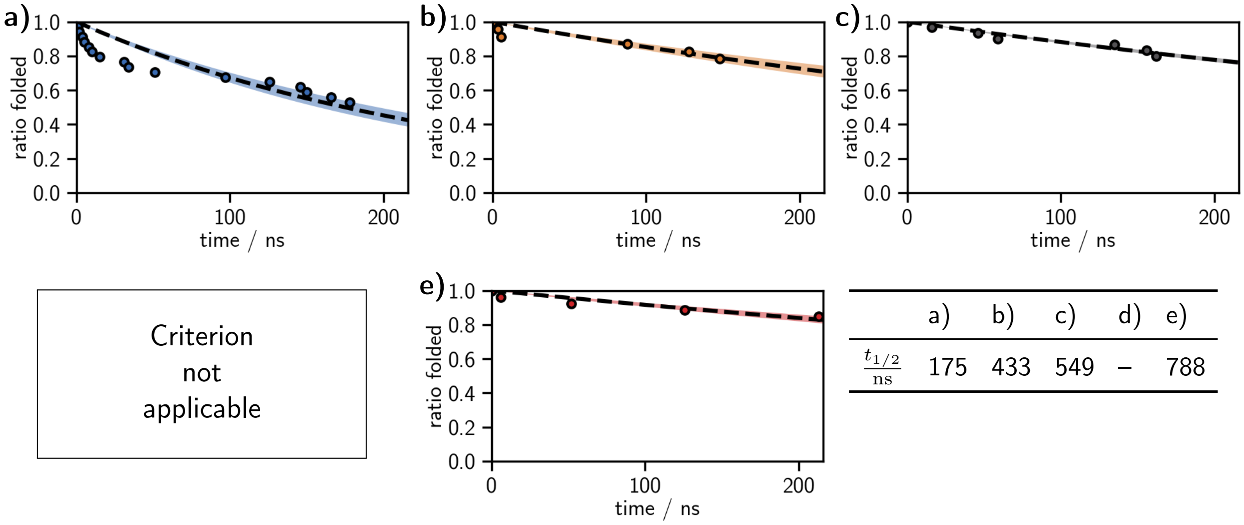


| | a) | b) | c) | d) | e) |
|---|---|---|---|---|---|
| $\frac{t_{1/2}}{\mathrm{ns}}$ | 175 | 433 | 549 | – | 788 |

Figure 25: Unfolding events in E285-protonated apo-langerin (a6/a7) and exponential fits measured by **a)** visual inspection (last folded frame), **b)** visual inspection (mean of last folded and first unfolded frame), **c)** RMSD cut-off (>0.2 nm, **d)** breaking of N287m–D308s H-bond (never observed), **e)** breaking of N288m–D308s H-bond.

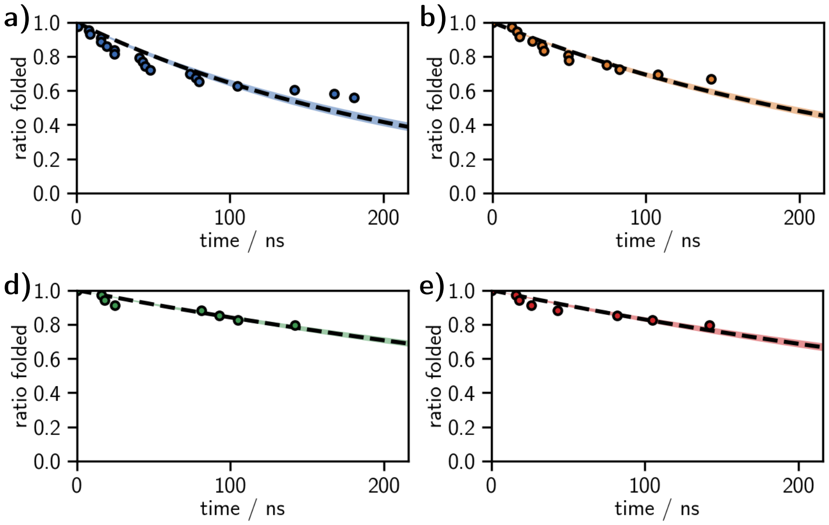


| | a) | b) | c) | d) | e) |
|---|---|---|---|---|---|
| $\frac{t_{1/2}}{\text{ns}}$ | 158 | 189 | 242 | 400 | 369 |

Figure 26: Unfolding events in E293-protonated apo-langerin (a10/a11) and exponential fits measured by **a)** visual inspection (last folded frame), **b)** visual inspection (mean of last folded and first unfolded frame), **c)** RMSD cut-off (>0.2 nm, **d)** breaking of N287m–D308s H-bond, **e)** breaking of N288m–D308s H-bond.

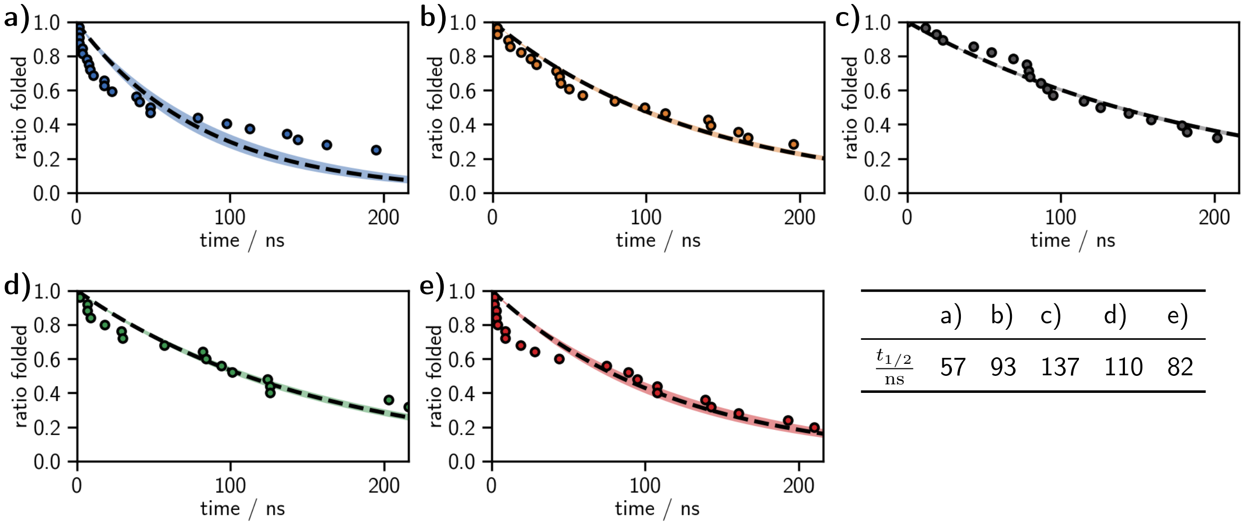


| | a) | b) | c) | d) | e) |
|---|---|---|---|---|---|
| $\frac{t_{1/2}}{\text{ns}}$ | 57 | 93 | 137 | 110 | 82 |

Figure 27: Unfolding events in D308-protonated apo-langerin (a18/a19) and exponential fits measured by **a)** visual inspection (last folded frame), **b)** visual inspection (mean of last folded and first unfolded frame), **c)** RMSD cut-off (>0.2 nm, **d)** breaking of N287m–D308s H-bond, **e)** breaking of N288m–D308s H-bond.

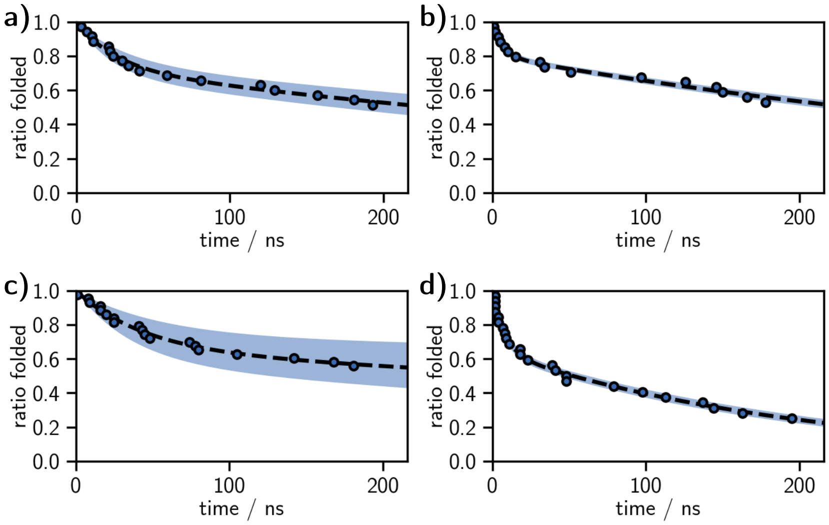


Figure 28: Unfolding events by visual inspection (last folded frame) and double exponential fits in **a)** neutral apo- (a0), **b)** E285 protonated (a6/a7), **c)** E293 protonated (a10/a11), and **d)** D308 protonated (a18/a19) langerin. The single exponential fit for the H294 protonated case is already sufficiently good (see fig. 24).

### ITC measurements

Data analysis, plotting and curve fitting was performed with OriginPro 2019 (OriginLab, Northampton, MA).

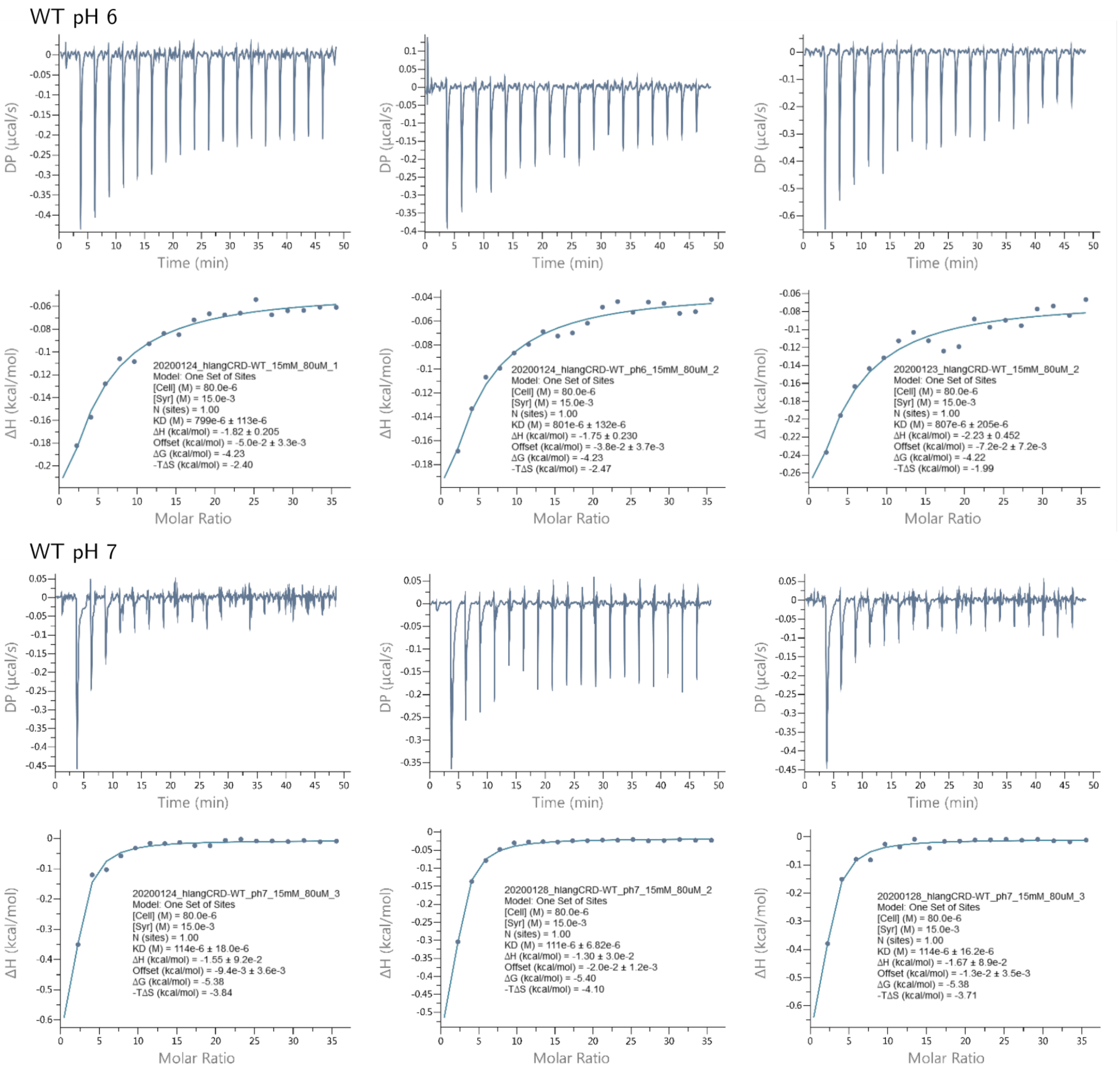


Figure 29: ITC measurements for wild type langerin.

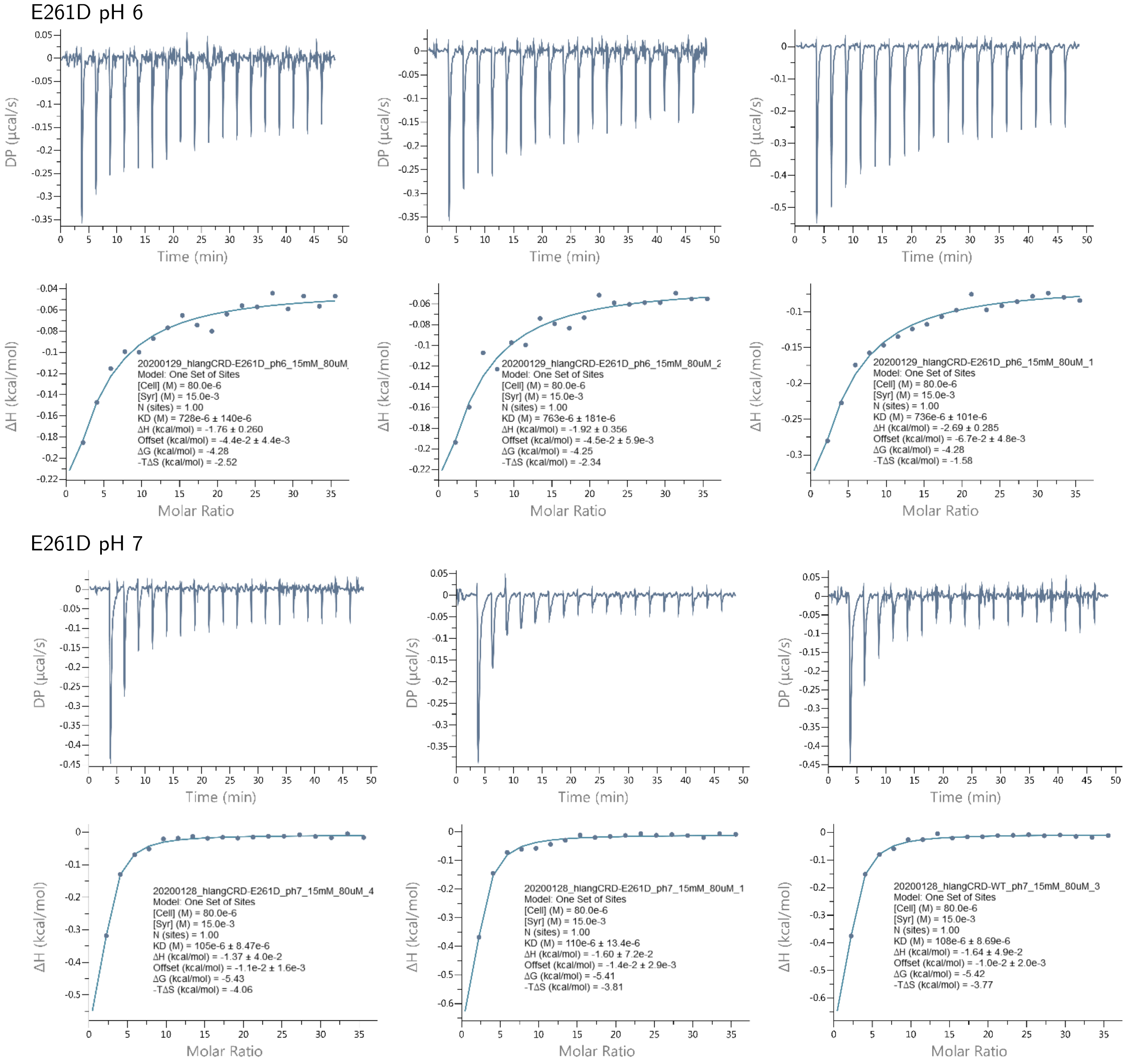


Figure 30: ITC measurements for the E261D langerin mutant.

#### Comparison of langerin to other C-type lectins

```
P07306|ASGR1_HUMAN|ASGPR    NWVEH------ERSCYWFSRSGKAW-ADADNYCRLEDAHLVVVTSWEEQKFVQHHIGPV-N--TWMGLHDQ--N 217
P16109|LYAM3_HUMAN|PSECtin  LISELTNQKEVAAWTYHYSTKAYSW-NISRKYCQNRYTDLVAIQNKNEIDYLNKVLPYY-SSYYWIGIRKN--N 98
P16581|LYAM2_HUMAN|ESECtin  ALTLVLLIKESGAWSYNTSTEAMTY-DEASAYCQQRYTHLVAIQNKEEIEYLNSILSYS-PSYYWIGIRKV--N 78
P14151|LYAM1_HUMAN|LSECtin  LCCDFLAHHGTDCWTYHYSEKPMNW-QRARRFCRDNYTDLVAIQNKAEIEYLEKTLPFS-RSYYWIGIRKI--G 95
P11226|MBL2_HUMAN|MBP       KWLTFSLGKQVGNKFFLTNGEIMTF-EKVKALCVKFQASVATPRNAAENGAIQNLI----KEEAFLGITDEKTE 172
Q6EIG7|CLC6A_HUMAN|Dectin2  SWKSF------GSSCYFISSEEKVW-SKSEQNCVEMGAHLVVFNTEAEQNFIVQQLNES-FS-YFLGLSDPQGN 146
Q9NNX6|CD209_HUMAN|DCSIGN   EWTFF------QGNCYFMSNSQRNW-HDSITACKEVGAQLVVIKSAEEQNFLQLQSSRS-NRFTWMGLSDLNQE 324
Q9H2X3|CLC4M_HUMAN|DCSIGNR  DWTFF------QGNCYFMSNSQRNW-HDSVTACQEVRAQLVVIKTAEEQNFLQLQTSRS-NRFSWMGLSDLNQE 336
Q9UJ71|CLC4K_HUMAN|Langerin GWKYF------KGNFYYFSLIPKTW-YSAEQFCVSRNSHLTSVTSESEQEFLYKTAGGL-I--YWIGLTKAGME 261
Q8VBX4|CLC4K_MOUSE|Langerin GWKYF------SGNFYYFSRTPKTW-YSAEQFCISRKAHLTSVSSESEQKFLYKAADGI-P--HWIGLTKAGSE 264
Q9ULY5|CLC4E_HUMAN|Mincle   NWEYF------QSSCYFFSTDTISW-ALSLKNCSAMGAHLVVINSQEEQEFLSYKKPKM-RE-FFIGLSDQVVE 147
Q8WXI8|CLC4D_HUMAN|MCL      DWRAF------QSNCYFPLTDNKTW-AESERNCSGMGAHLMTISTEAEQNFIIQFLDRR-LS-YFLGLRDENAK 151
P35247|SFTPD_HUMAN|SP       KVELFPNGQSVGEKIFKTAGFVKPF-TEAQLLCTQAGGQLASPRSAAENAALQQLVVAK-NEAAFLSMTDSKTE 321
Q5KU26|COL12_HUMAN|SR       HWKNF------TDKCYYFSVEKEIF-EDAKLFCEDKSSHLVFINTREEQQWIKK-QMVG-RESHWIGLTDSERE 674
P06734|FCER2_HUMAN|CD23     KWINF------QRKCYYFGKGTKQW-VHARYACDDMEGQLVSIHSPEEQDFLTKHASHT-G--SWIGLRNLDLK 229
Q9UBG0|MRC2_HUMAN|Endo180   SWQPF------QGHCYRLQAEKRSW-QESKKACLRGGGDLVSIHSMAELEFITKQIKQE-VEELWIGLNDLKLQ 449
P22897|MRC1_HUMAN|MMR       QWWPY------AGHCYKIHRDEKKIQRDALTTCRKEGGDLTSIHTIEELDFIISQLGYEPNDELWIGLNDIKIQ 432
                                           :                *      :    .  *   :            ::.: .

P07306|ASGR1_HUMAN|ASGPR    GPWKWVDGTDYETGF--KNWRPEQPDDWYGHGLGGGEDCAHF-----TDDGRWNDDVCQRPY-RWVCETELDK 282
P16109|LYAM3_HUMAN|PSECtin  KTWTWVGTKKALT-NEAENWADNEPNNK-----RNNEDCVEIYIKSPSAPGKWNDEHCLKKK-HALCYTASCQ 164
P16581|LYAM2_HUMAN|ESECtin  NVWVWVGTQKPLT-EEAKNWAPGEPNNR-----QKDEDCVEIYIKREKDVGMWNDERCSKKK-LALCYTAACT 144
P14151|LYAM1_HUMAN|LSECtin  GIWTWVGTNKSLT-EEAENWGDGEPNNK-----KNKEDCVEIYIKRNKDAGKWNDDACHKLK-AALCYTASCQ 161
P11226|MBL2_HUMAN|MBP       GQFVDLTGNRLTY----TNWNEGEPNNA-----GSDEDCVLL-----LKNGQWNDVPCSTSH-LAVCEFPI-- 228
Q6EIG7|CLC6A_HUMAN|Dectin2  NNWQWIDKTPYEKNV--RFWHLGEPNH-----SA--EQCASIVFW-KPTGWGWNDVICETRR-NSICEMNKIY 208
Q9NNX6|CD209_HUMAN|DCSIGN   GTWQWVDGSPLLPSF-KQYWNRGEPNN-----VGE-EDCAE------FSGNGWNDDKCNLAK-FWICKKSAAS 381
Q9H2X3|CLC4M_HUMAN|DCSIGNR  GTWQWVDGSPLSPSF-QRYWNSGEPNN-----SGN-EDCAE------FSGSGWNDNRCDVDN-YWICKKPAA- 394
Q9UJ71|CLC4K_HUMAN|Langerin GDWSWVDDTPFNKVQSVRFWIPGEPNN-----AGNNEHCGNIKA---PSLQAWNDAPCDKTF-LFICKRPYVP 325
Q8VBX4|CLC4K_MOUSE|Langerin GDWYWVDQTSFNKEQSRRFWIPGEPNN-----AGNNEHCANIRV---SALKCWNDGPCDNTF-LFICKRPYVQ 328
Q9ULY5|CLC4E_HUMAN|Mincle   GQWQWVDGTPLTKSL--SFWDVGEPNN-----IATLEDCATMRDS-SNPRQNWNDVTCFLNY-FRICEMVGIN 211
Q8WXI8|CLC4D_HUMAN|MCL      GQWRWVDQTPFNPRR--VFWHKNEPDN-----SQ--GENCVVLVY-NQDKWAWNDVPCNFEA-SRICKIPGTT 213
P35247|SFTPD_HUMAN|SP       GKFTYPTGESLVY----SNWAPGEPNDD-----GGSEDCVEI-----FTNGKWNDRACGEKR-LVVCEF---- 375
Q5KU26|COL12_HUMAN|SR       NEWKWLDGTSPDY----KNWKAGQPDNW-GHGHGPGEDCAGL-----IYAGQWNDFQCEDVN-NFICEKDRET 736
P06734|FCER2_HUMAN|CD23     GEFIWVDGSHVDY----SNWAPGEPTS-----RSQGEDCVMM-----RGSGRWNDAFCDRKLGAWVCDRLATC 288
Q9UBG0|MRC2_HUMAN|Endo180   MNFEWSDGSLVSF----THWHPFEPNNF----RDSLEDCVTIW----GPEGRWNDSPCNQSL-PSICKKAGQL 510
P22897|MRC1_HUMAN|MMR       MYFEWSDGTPVTF----TKWLRGEPSHE----NNRQEDCVVMK----GKDGYWADRGCEWPL-GYICKMKSRS 492
                              :                *   :*            .              * *  *       :*
```

**Legend**

Uniprot annotations:
- * fully conserved
- : strongly similar
- . weakly similar

Short-loop
Long-loop

- **W** Conserved residue found in langerin
- H HIS294
- H Other histidines in interesting positions
- E Primary $Ca^{2+}$ binding-site residue
- E Secondary $Ca^{2+}$ binding-site residue
- K K257 or analogue
- K K close to K257 analogue

Figure 31: Sequence alignment using UniProtKB (27) for langerin's carbohydrate recognition domain (residues 198 to 325) with selected C-type lectins. Annotations have been added by comparing available crystal structures of ASGPR (PDB 5JPV), PSECtin (PDB 1G1S), ESECtin (PDB 1G1T), LSECtin (PDB 5VC1), MBP (PDB 1HUP), dectin-2 (PDB 5VYB), DC-SIGN (PDB 1SL4), DC-SIGNR (PDB 1XPH), Mincle (PDB 3WH2), MCL (PDB 3WHD), SP (PDB 1PWD), SR (PDB 2OX8), CD23a (PDB 4G9A), Endo180 (PDB 5AO5), and MMR (PDB 5XTS).

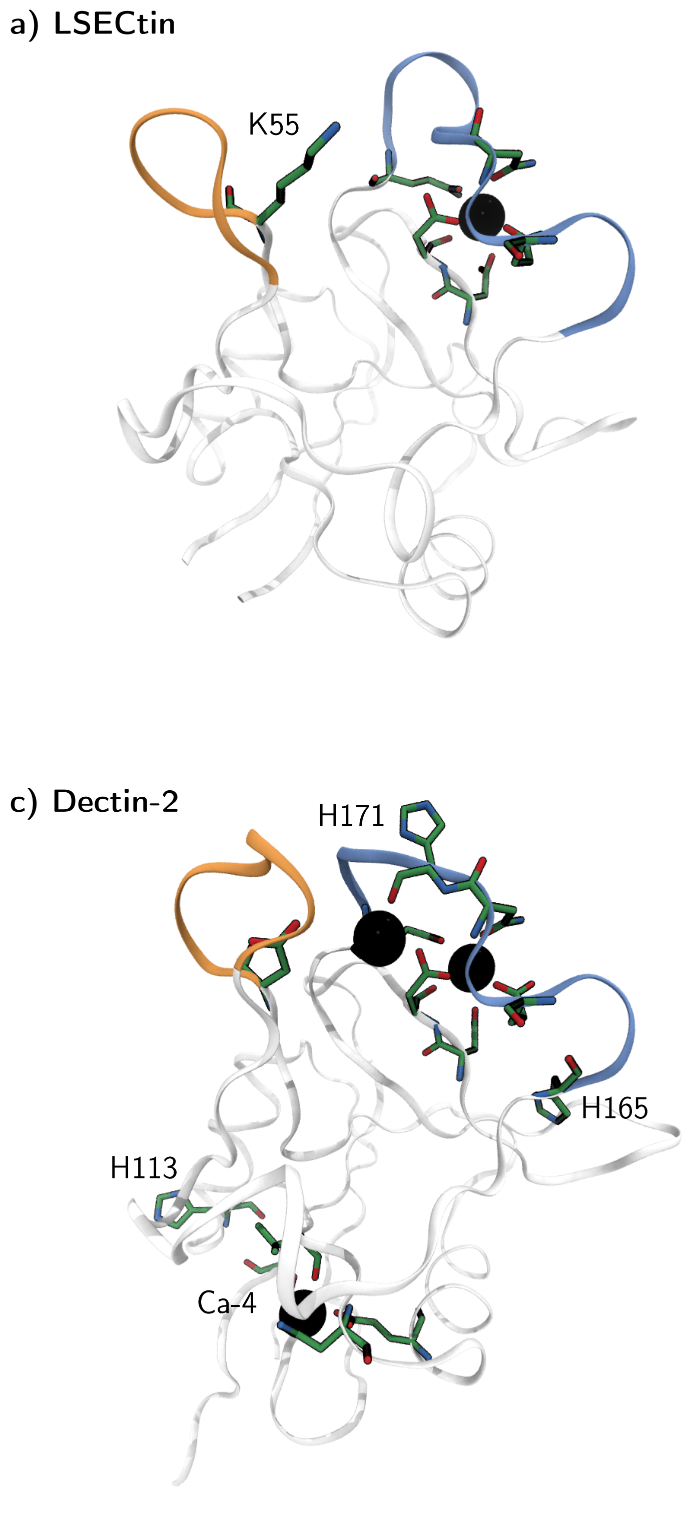


Figure 32: **a)** LSECtin (PDB 5VC1) has a lysine residue (K55) at the same position as langerin (K257). **b)** The carbohydrate recognition domains of C-type lectins may contain multiple secondary $Ca^{2+}$ binding-sites (Ca-1, Ca-3) like it is the case for DC-SIGN (PDB 1SL4). **c)** Potentially pH-sensitive histidine residues are found for example in dectin-2 (PDB 5VYB) in two positions close to the primary $Ca^{2+}$ binding site. A moderately conserved histidine H113 (H229 in langerin) can be found close to a secondary bindig-site (Ca-4). **d)** ASGPR (PDB 5JPV) with pH sensing residue H256. The distance between the histidine side-chain and the $Ca^{2+}$-ion in the binding pocket is about 0.6 nm in this structure and potentially lower in the protonated state due to a likely interaction with D266. This close proximity suggests a pH-switching mechanism for the $Ca^{2+}$-affinity through direct interaction.
